# Single-cell T cell receptor sequencing deciphers αβ- and γδ-T cell responses in neuroblastoma

**DOI:** 10.64898/2026.11.18.767512

**Authors:** Elisa Zappa, Naima Hiddink Verberne, Stijn A. Messing, Fajar Purayamkott Murikkumchery, Guillem Sánchez Sánchez, Sofia Almeida Andrezo, Stijn De Munter, Bart Vandekerckhove, Chin Leng Tan, Aleksandra K. Balwierz, Wim J. de Jonge, Jeff DeMartino, Philip Lijnzaad, Thanasis Margaritis, Anne Laura Borst, Alida F. W. van der Steeg, Ronald de Krijger, Miranda P. Dierselhuis, Godelieve A. M. Tytgat, Monika C. Wolkers, Edward W. Green, Wouter Scheper, David Vermijlen, Jan J. Molenaar, Judith Wienke

**Affiliations:** Princess Máxima Center for Pediatric Oncology, Utrecht, the Netherlands; Institute of Medical Immunology, Université Libre de Bruxelles (ULB), Gosselies, Belgium; ULB Center for Research in Immunology (U-CRI), ULB, Brussels, Belgium; Department of Pharmacotherapy and Pharmaceutics, ULB, Brussels, Belgium; WELBIO Department, WEL Research Institute, Wavre, Belgium; Department of Diagnostic Sciences, Ghent University, Ghent, Belgium; Cancer Research Institute Ghent (CRIG), Ghent, Belgium; GMP Unit CellGenTherapies, Ghent University Hospital, Ghent, Belgium; German Cancer Research Center (DKFZ) Heidelberg, CCU Neuroimmunology and Brain Tumor Immunology, Heidelberg, Germany; German Cancer Consortium (DKTK), DKFZ, Core Center Heidelberg, Heidelberg, Germany; Department of Neurology, Medical Faculty Mannheim, MCTN, University of Heidelberg, Mannheim, Germany; DKFZ Hector Cancer Institute at the University Medical Center, Mannheim, Germany; Oncode Institute, Utrecht, the Netherlands; Department of Pathology, University Medical Center Utrecht, Utrecht, The Netherlands; Sanquin Blood Supply Foundation, Department of Research, T cell differentiation lab; Amsterdam UMC, University of Amsterdam, Landsteiner Laboratory, Amsterdam, The Netherlands; Division of Molecular Oncology & Immunology, The Netherlands Cancer Institute, Amsterdam, the Netherlands

**Author notes:** These authors contributed equally (Elisa Zappa, Naima Hiddink Verberne, David Vermijlen, Jan J. Molenaar).

## Abstract

High-risk neuroblastoma is a pediatric tumor with poor survival rates and low MHC-I expression. The cellular determinants of anti-tumor immunity remain poorly defined, limiting development of targeted immunotherapies. We generated a single-cell atlas integrating transcriptomic and αβ/γδ T cell receptor (TCR) sequencing of >150,000 T cells from tumor and blood of 10 patients throughout treatment. Fractions of CD8^+^ and CD4^+^ T cells are clonally expanded and activated, with tumor-specific expansion among CD4^+^ cells. Transcriptomic evidence of tumor-reactivity is more pronounced in CD4^+^ than CD8^+^ T cells. Moreover, Vδ1^+^ γδT cells are enriched, displaying TCR engagement. Functional screening confirmed reactivity of selected γδTCRs and, remarkably, a small fraction of CD8^+^ αβTCRs. Subset-specific inhibitory receptors, including CTLA4, FCRL6, and KIRs, may serve as targets for tailored immunotherapies. Together, we identify distinct αβ- and γδT cell responses despite low MHC-I, and provide a comprehensive resource for deciphering pediatric immunity against MHC-I^low^ tumors.

## Introduction

T cell-based immunotherapies have transformed cancer treatment, demonstrating remarkable efficacy in hematological malignancies and immunologically responsive solid tumors like melanoma^1–3^. Yet, their therapeutic potential in pediatric solid tumors remains largely unrealized, with high-risk neuroblastoma representing a particularly challenging tumor with survival rates <60%^4,5^. Despite intensive multimodal treatment, including chemotherapy, radiotherapy, and immunotherapy with anti-GD2 antibodies, many patients experience disease relapse and ultimately succumb to their disease^4^. Although emerging T cell-based approaches, including chimeric antigen receptor (CAR) T cell therapy, have shown encouraging clinical activity in neuroblastoma, durable responses remain limited, with early relapse and treatment failure particularly in patients with high tumor burden^5^. These observations highlight the urgent need for more effective immunotherapeutic strategies.

A deeper understanding of the endogenous anti-tumor immune response is currently lacking and will be essential to identify key determinants of effective tumor control. A major challenge for T cell-mediated tumor recognition in neuroblastoma is its characteristically low expression of major histocompatibility complex class I (MHC-I), which limits the presentation of tumor-derived peptides to conventional CD8⁺ αβT cells^6^. This is particularly relevant for endogenous T cell responses, as native TCR-mediated recognition depends on the availability of cognate antigen presented by the appropriate MHC molecules. In addition, the low tumor mutational burden of neuroblastoma limits the generation of tumor-specific neoantigens^7^. Nevertheless, the positive association between intratumoral T cell infiltration and patient survival suggests that T cells relevantly contribute to tumor control^6,8^. To date, the cellular and molecular determinants of anti-tumor immunity in neuroblastoma remain however poorly defined. For instance, despite expression of the inhibitory ligand PD-L1 in neuroblastoma^9^, immune checkpoint blockade targeting the PD-1 axis has shown limited clinical efficacy^10^. This may reflect lacking endogenous tumor reactivity, or additional immunosuppressive mechanisms within the tumor microenvironment, including inhibitory pathways involving TIGIT and macrophage migration inhibitory factor (MIF), for which we and others have previously reported evidence^11,12^. Together, these observations raise a fundamental question: whether conventional αβT cell responses are mounted against neuroblastoma and, more broadly, which T cell populations contribute to tumor recognition in the context of low MHC-I expression.

Beyond conventional CD8^+^ αβT cells, alternative T cell compartments may contribute to anti-tumor immunity. γδT cells are particularly compelling candidates, as their recognition of target cells can occur independently of classical peptide-MHC presentation, providing a potential route for tumor recognition in MHC-I-low settings^13^. In other malignancies the contribution of γδT cells to immune surveillance and tumor control, particularly in tumors with impaired classical MHC expression, has already been established^14,15^. In parallel, CD4^⁺^ αβT cells may contribute to tumor immunity through both direct and indirect mechanisms, including tumor-cell recognition in the context of MHC-II presentation and the orchestration of broader anti-tumor immune responses^16,17^. Thus, distinct αβ- and γδT cell populations represent potentially important, yet incompletely characterized components of the immune response against neuroblastoma. Identifying tumor-reactive populations and intrinsic or extrinsic mechanisms that constrain their activity will be essential to identify actionable mechanisms that may be exploited to enhance anti-tumor immunity.

Here, we generated a comprehensive single-cell T cell atlas of neuroblastoma by integrating single-cell transcriptomic profiling with αβ/γδTCR sequencing of tumor and blood samples of 10 patients throughout treatment. This integrated approach enabled the characterization of T cell subset composition, clonal dynamics, transcriptional states, potential tumor reactivity, and targetable inhibitory receptors across distinct αβ- and γδT cell subsets. Moreover, functional screening confirmed the reactivity of selected CD8^⁺^ αβ- and γδTCRs. Together, our findings provide a deep characterization of T cell responses against neuroblastoma, revealing distinct patterns of αβ- and γδT cell activation and tumor reactivity. This dataset provides a comprehensive resource for deciphering pediatric immunity against MHC-I^low^ tumors and a framework for the development of rationally designed T cell-based immunotherapies.

## Results

### I. Neuroblastoma patients exhibit enhanced T cell memory differentiation profiles

To characterize the T cell dynamics in neuroblastoma patients throughout chemotherapy, we enriched paired tumor and blood samples of 10 high-risk neuroblastoma (HR-NB) patients for CD3^+^ T cells while also maintaining other cell types (∼75% CD3^+^ and ∼25% other cells; see method section for details). The selected cells were subjected to parallel single-cell RNA, αβTCR and γδTCR sequencing (scRNAseq, scTCRseq). Pediatric age-matched healthy donor (HD) blood samples were used as controls^18^ (**Figure 1A**). This yielded a comprehensive T cell-rich transcriptome atlas comprising 11’558 tumor cells, 21’059 B cells, 23’315 myeloid cells 44’045 NK cells, and 180’212 T cells (**Figure S1A-D**).

**Figure 1:**
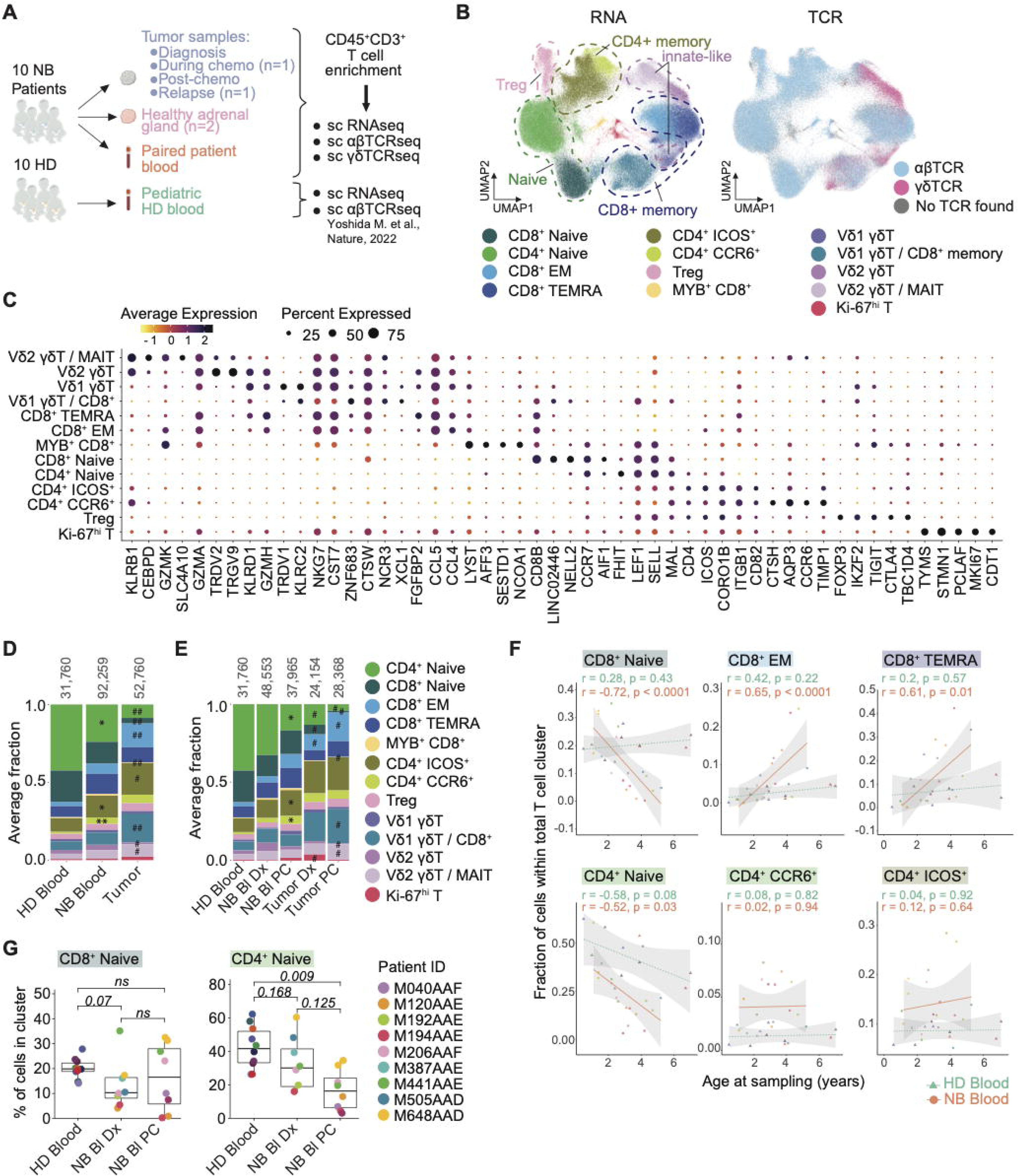
A premature tumor-driven differentiated phenotype is observed in NB circulating T cells. **(A)** Schematic representation of the study. NB: neuroblastoma; HD: healthy donor; sc: single-cell; TCR: T cell receptor. **(B)** UMAP dimensionality reduction of T of patients and HD colored by annotation and by αβTCR and gdTCR expression **(C)** Expression of the top 5 genes per each cluster of figure 1B. Color represents average expression. Size of the dot represents percentage of cells expressing the marker. **(D)** Average cluster frequency per patient/HD samples within HD blood, NB blood and tumor. Numbers on top of the bar indicate the total cells per sample. * = statistical difference between HD and NB blood; # = statistical difference between NB blood and tumor. Only statistically significant comparisons are depicted. **(E)** Average cluster frequency per patient/HD samples within HD blood, NB blood and tumor pre- and post-chemo. Numbers on top of the bar indicate the total cells per sample. * = statistical difference between HD and NB blood; # = statistical difference between NB blood and tumor. Only statistically significant comparisons are depicted. *Bl: Blood; Dx: diagnosis; PC: post-chemo.* **(F)** Percentage of CD8^+^ naïve, CD4^+^ naïve, CD8^+^ EM, CD8^+^ TEMRA, CD4^+^ ICOS^+^ and CD4^+^ CCR6^+^ T cells in HD (green, dashed) and NB (red, full) blood at different time of patient age at sampling (years). Each dot/triangle represents a patient/HD. **(G)** Percentage of CD8^+^ naïve and CD4^+^ naïve cells per patient/HD sample within HD blood, NB blood at diagnosis, NB blood post-chemo. Each dot represents a patient/HD. Samples from the same patient pre- and pos-chemo re indicated with the same color. *Bl: Blood; Dx: diagnosis; PC: post-chemo*.

Within the T cell population, we identified 10 clusters of αβT cells and 4 clusters of unconventional (including γδ) T cells (**Figure 1B-C**). Within CD4^+^ αβT cells, we identified CD4^+^ naïve cells (CCR7^+^ SELL^+^), two clusters of memory CD4^+^ T cells (ICOS^+^ c.q. CCR6^+^) and regulatory T cells (Tregs; FOXP3^+^CTLA4^+^TIGIT^+^). The CD8^+^ αβT cell population comprised CD8^+^ naive cells (CCR7^+^ SELL^+^), CD8^+^ effector memory (EM) cells (GZMK^+^ HLA-DRB1^+^ GZMH^+^ GZMA^+^), CD8^+^ terminally differentiated effector memory (TEMRA) cells (GZMB^+^ HLA-DRB1^+^ GZMH^+^ GZMA^+^), and a cluster composed of mixed populations of αβ CD8^+^ memory T cells and Vδ1^+^ γδT cells, a subset of γδT cells characterized by usage of the TRDV1 chain^19^. In addition, we identified a separate TRDV1^+^ and a Vδ2^+^ (TRDV2^+^) γδT cell cluster and a few smaller clusters (proliferating MKI67^+^ (Ki-67^+^) T cells and MYB^+^ CD8^+^ T cells), with ≤3,000 cells each (**Figure 1C, S2A-B**).

To evaluate compositional differences between pediatric HD and patient samples, we compared the frequencies of the distinct T cell clusters. In accordance with previously published observations of memory skewing in tissue/tumor-infiltrating T cells^20^, we observed decreased frequencies of CD4^+^ and CD8^+^ naive T cells in the tumor compared to patient and HD blood (**Figure 1D, S2C**). Strikingly, frequencies of naive αβT cells were not only reduced in the tumor, but also in patient blood compared to pediatric HD blood, while memory T cell frequencies were increased, particularly those of CD4^+^ T cells (**Figure 1D**). Moreover, a naive-memory shift was observed between pre- and post-chemotherapy samples (**Figure 1E**). In line with this finding, the frequency of naive CD4^+^ T cells declined over time, yet both in patient and HD blood (**Figure 1F**). More strikingly, we observed a steep decline in the frequency of naive CD8^+^ T cells and an increase in memory CD8^+^ T cells in the blood with patient age – a feature that we did not observe in HD peripheral T cells (**Figure 1F**). This finding suggests that CD8^+^ T cells undergo enhanced differentiation in patients both in the tumor and in blood, similar to previous reports in adult follicular B cell lymphoma patients^21^. Of note, the reduced frequency of circulating CD8^+^ naive T cells was already observed at diagnosis, before chemotherapy (**Figure 1G**). These results indicate that tumor presence may drive accelerated memory-differentiation of αβT cells in neuroblastoma patients.

### II. Fractions of memory αβT cells in neuroblastoma tumors are clonally expanded

We reasoned that αβT cell antigen encounter may drive the observed accelerated naive-to-memory conversion. To test this hypothesis, we evaluated the clonal expansion of αβT cell clones in tumor samples, NB blood and in HD blood, with the assumption that T cell triggering results in clonal expansion, which can therefore be used as a surrogate for antigen-experienced T cells. We defined a clone as expanded when we identified ≥2 TCR copies within the same sample (e.g.: tumor at diagnosis for a given patient). >95% of clonally expanded αβTCRs were not found in datasets of known virus-specific TCRs, suggesting that the bulk of our identified TCRs are unlikely to be virus-reactive (**Figure S3A-B**). A higher fraction of αβT cells was clonally expanded in patient blood than HD blood, and the highest clonality was observed in the tumor (**Figure 2A-B**). Yet, clonal expansion rates displayed high sample heterogeneity, ranging from 0-60% clonally expanded fractions per sample (**Figure 2A-B**). Expansion rates of αβT cells did not significantly differ before and after chemotherapy, suggesting a limited effect of chemotherapy on T cell clonality (**Figure 2B-C**).

**Figure 2:**
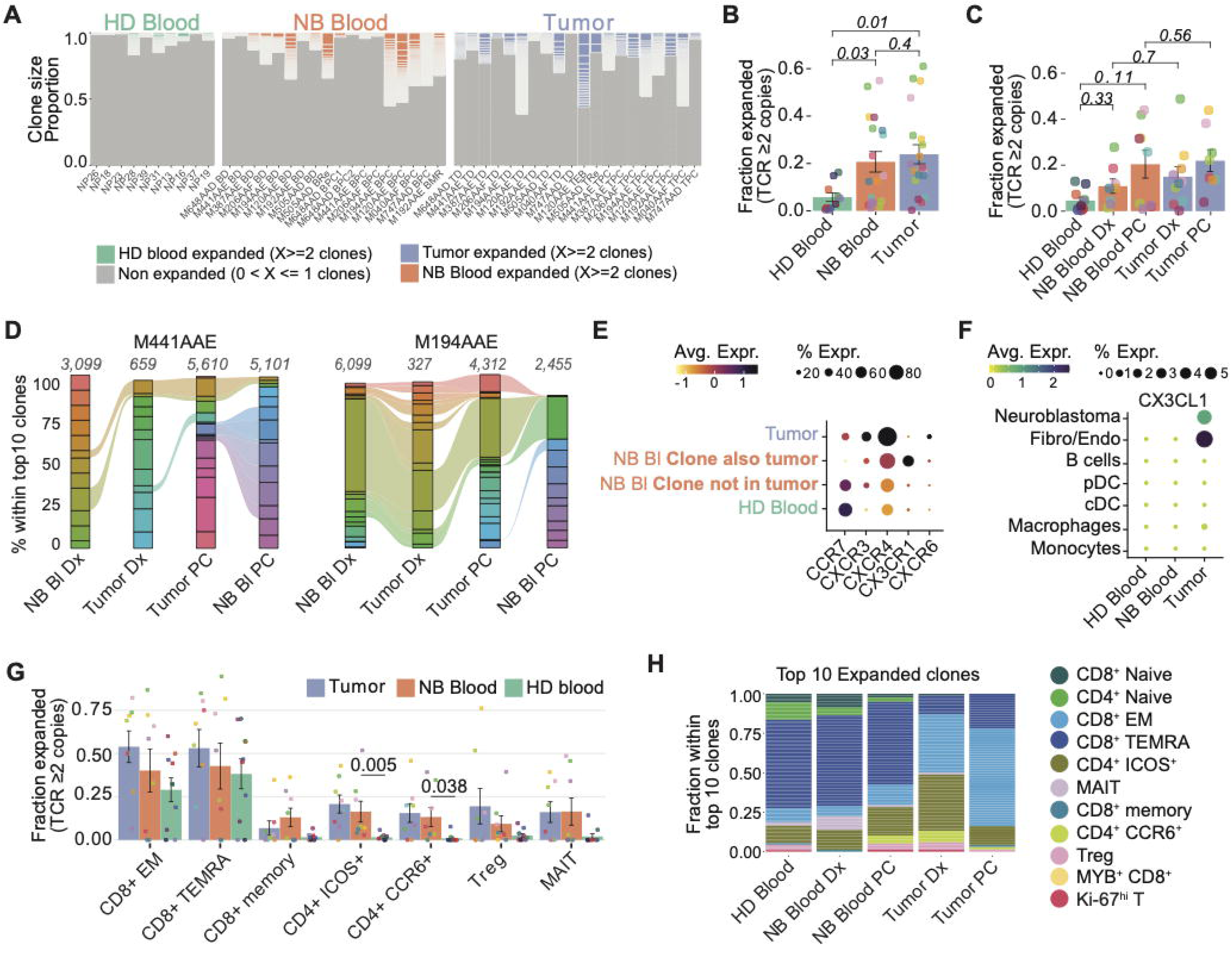
Memory αβT cells are expanded and recirculate. **(A)** Clone size proportion of single (grey) and expanded (colored) αβT cell clones within each HD blood (green), NB blood (red) and tumor (blue) sample. **(B)** Average expansion proportion of αβ T cell clones within HD blood (green), NB blood (red) and tumor (blue). **(C)** Average expansion proportion of αβ T cell clones within HD blood (green), NB blood (red) and tumor (blue) pre- and post-chemotherapy. Dx: diagnosis; PC: post-chemotherapy. **(D)** Proportion of top 10* αβT cell clonotypes in the repertoire normalized to 100% in blood and tumor at different therapy timepoints, for two representative patients. * Indicates that bars over 100% contain clones not within the top 10 of a specific sample. Numbers on top of the bar indicate total number of αβT cell clones within sample. Dx: diagnosis; PC: post-chemotherapy. **(E)** Expression of chemokine receptors by HD blood, NB circulating αβT cells present/absent (recirculating/non-recirculating) in tumor. Color represents average expression. Size of the dot represents percentage of cells expressing the marker. Bl: blood. **(F)** Expression of CX3CR1 ligand, CX3CL1 in the microenvironment. Color represents average expression. Size of the dot represents percentage of cells expressing the marker. **(G)** Average proportion per sample of expanded αβT cell clones in HD blood (green), NB blood (red) post-chemotherapy and tumor (blue) post-chemotherapy. **(H)** Top 10 expanded αβT cell clone subset composition within HD blood (green), NB blood (red) and tumor (blue) pre- and post-chemo colored by annotation. Dx: diagnosis; PC: post-chemotherapy.

To evaluate the TCR repertoire evolution throughout chemotherapy, we followed the top 10 expanded clonotypes for each patient within blood and tumor, pre- and post-treatment. A proportion of the top 10 T cell clones present in the tumor at diagnosis persisted as expanded clones during chemotherapy, suggesting a robust, prolonged persistence. The transcriptomic profiles of these persistent clones did not markedly differ before and after chemotherapy but showed some enrichment of pathways associated with SLIT/ROBO regulation and nonsense-mediated decay post-chemotherapy **(Figure S3C-D)**. Despite the high variability observed in clonal expansion rates, for each patient one or more of these expanded tumor-infiltrating clones was also detected in matched blood, suggestive of migration between tissue compartments (**Figure 2D** and **Figure S3E-F**). To support this notion, we analyzed the chemokine receptor profile of T cell clones which were either exclusively detected in blood or tumor, or found in both blood and tumor (“migrating”). We selected for chemokine receptors with expression by at least 20% of cells. As expected, CCR7 was higher in blood-derived T cells, and CXCR3/4/6 were higher in tumor-derived T cells. Furthermore, we observed expression of CX3CR1 specifically on T cells from blood whose TCR clonotypes were also detected in the tumor (**Figure 2E**). CX3CR1 has been previously reported to drive recruitment of circulating T cells to tumor sites^22^. Furthermore, its ligand CX3CL1 was selectively expressed by neuroblastoma and fibroblasts/endothelial cells in the TME (**Figure 2F**), indicative of recruitment of circulating clones to the tumor through the CX3CL1-CX3CR1 axis.

We next investigated whether the clonal expansion profiles in distinct tissue compartments differed between T cell subsets. The highest clonal expansion rates in the tumor were observed in the CD8^+^ TEMRA and CD8^+^ EM compartments, but expansion rates did not significantly differ from NB blood and HD blood, suggesting limited tumor-specific, locally enhanced clonal expansion (**Figure 2G, S3F-G**). The top 10 most clonally expanded T cell clones per patient were dominated by CD8^+^ TEMRA in blood, and by CD8^+^ EM cells in the tumor (**Figure 2H**). Intriguingly, patient-specific and tumor-specific expansion was rather observed in CD4^+^CCR6^+^ and CD4^+^ICOS^+^ T cell memory subsets (**Figure 2G, S3G-H**). The fraction of CD4^+^ T cells among the top 10 most expanded clones was higher in tumors at diagnosis than post-chemotherapy, suggesting a shift in TIL subset responses from CD4^+^ to CD8^+^ T cells during chemotherapy (**Figure 2H, S3H**). Overall, we found evidence of long-term persistent, clonally expanded αβT cells within patient tumors, specifically in the CD8^+^ T cell compartment, while tumor-specific clonal expansion was observed rather among CD4^+^ T cells.

### III. CD8^+^ αβ T cells are activated without evidence of persistent tumor-reactivity

Since a direct correlation between clonal expansion and tumor responsiveness has been described in adult cancers^23^, we assessed the transcriptomic profile of CD4^+^ and CD8^+^ memory T cell subsets within the tumor bed. CD4^+^CCR6^+^ T cells expressed markers indicative of a Th17 differentiation profile (CCR6^+^ IL17A^+^ RORC^+^ IL1R1^+^), which has been previously associated with a tumor-promoting phenotype, although highly context-dependent^24^. CD4^+^ICOS^+^ were positive for T follicular helper cell (Tfh)-associated markers (CXCL13^+^ CXCR5^+^ IL21^+^ TOX2^+^), which indicate an anti-tumorigenic role of the ICOS^+^ CD4^+^ subset via B-cell help, (im)mature tertiary lymphoid structure (TLS) formation and CD8^⁺^ T cell recruitment (**Figure 3A, Table S1)**^25^. (Im)mature disorganized TLS have been recently described particularly in high-risk neuroblastoma^26^. Indeed, we observed the presence of CXCR5^+^ B cells and BCL6^+^ dendritic cells within tumors, key subsets for TLS formation **(Figure S4A)** and a predicted TLS-supporting CD40-CD40L interaction between ICOS^+^CD4^+^ memory T cells and B cells (**Figure S4B)**. This suggests a role for CD4^+^ICOS^+^ T cells in the broader orchestration of neuroblastoma anti-tumor immunity.

**Figure 3:**
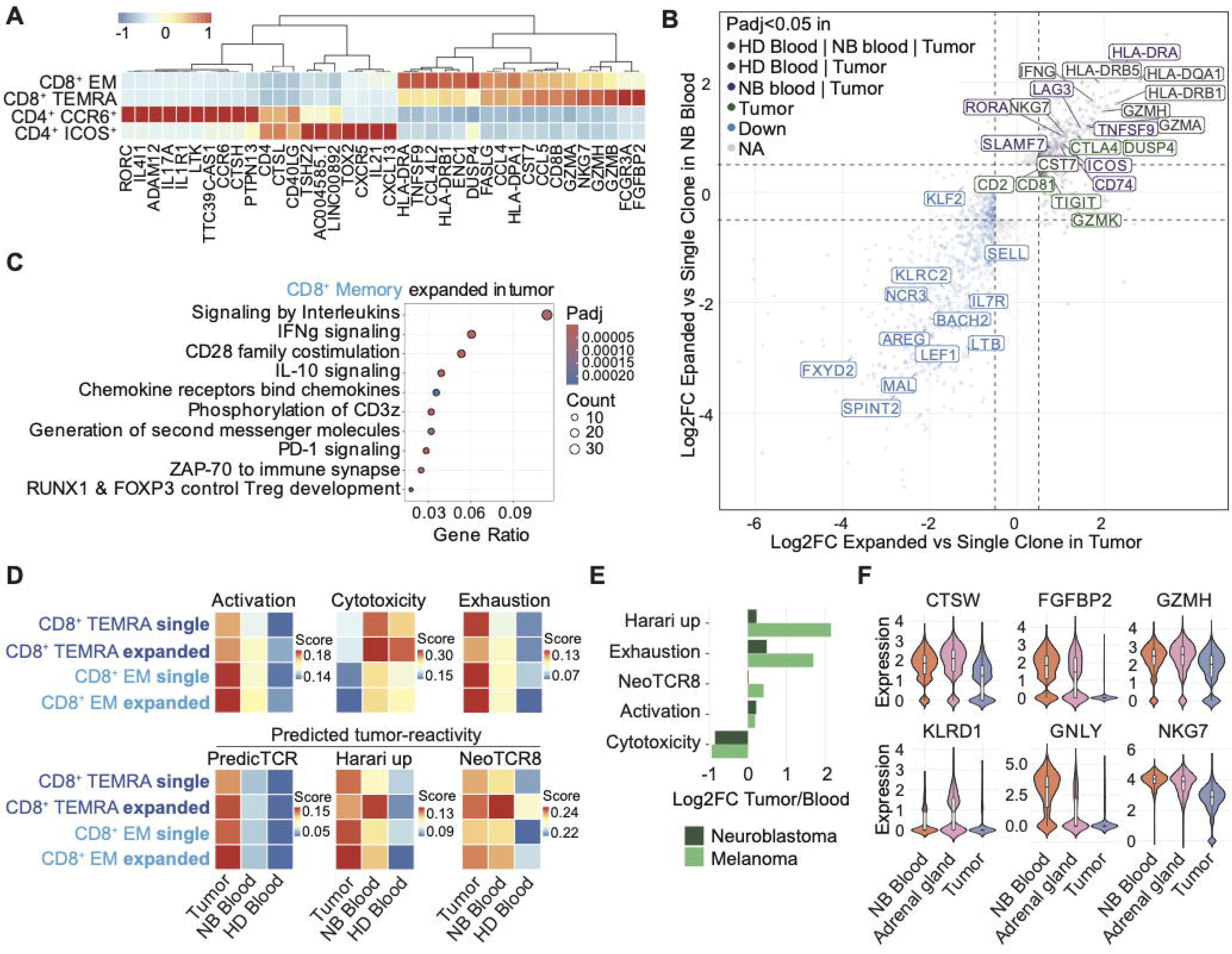
CD8+ memory T cells show reduced cytotoxicity in the tumor. **(A)** Expression of top10 markers for CD8^+^ EM, CD8^+^ TEMRA, CD4^+^ ICOS^+^ and CD4^+^ CCR6^+^ within the tumor bed. **(B)** Crater plot of significant differentially expressed genes of tumor expanded vs single clones within the CD8^+^ memory subsets. Average log2 foldchange in tumor and blood samples of these genes is shown. Green indicates genes upregulated only in expanded clones in the tumor; lilac indicates genes upregulated in expanded clones both in tumor and NB blood. **(C)** Pathway analysis showing Reactome pathways enriched in tumor expanded clones within the CD8^+^ memory. **(D)** Average value of activation, cytotoxicity, exhaustion and predicted tumor reactivity (by predicTCR, Harari_up and NeoTCR8 scores) within single and expanded CD8^+^ memory subsets in tumor, NB and HD blood. **(E)** Log2 Foldchange of UCell scores for activation, cytotoxicity, exhaustion and predicted tumor reactivity (by Harari_up and NeoTCR8 scores) in tumor versus blood of CD8^+^ memory T cells in our neuroblastoma dataset and in a public melanoma dataset. **(F)** Expression of cytotoxicity-related genes in the CD8^+^ memory T cells in neuroblastoma patient blood, tumor and matched normal adrenal gland samples. Statistics not shown due to sample numbers (n=2 samples).

Tumor-infiltrating CD8^+^ memory T cells displayed an activated phenotype (HLA-DP/DR^+^ GZMA/B/H^+^ FASLG^+^) (**Figure 3A, Table S1)**, in line with the observed accelerated peripheral CD8^+^ naive-to-memory conversion and overall high rates of clonal expansion in patients (**Figure 1F and Figure 2G-H**). However, neuroblastoma tumors express low levels of MHC-I molecules **(Figure S4C-D)**, posing a barrier for recognition by CD8^+^ T cells. This raises the question whether tumor-infiltrating CD8^+^ T cells may be tumor reactive or rather bystander T cells.

To investigate this, we performed differential gene expression analysis between expanded and non-expanded T clones, in blood versus tumor **(Figure 3B, Figure S4E, Table S2-4**). Overall, we observed a high correlation between expansion-specific genes in tumor, NB blood and HD blood, pointing towards a general activated, cytotoxic profile (HLA-DR, NKG7, GZMA, GZMH, CCL4) of tumor-infiltrating CD8+ memory T cells **(Figure 3B, S4F-G)**. Only few genes were specifically upregulated in patient samples (blood & tumor) compared to HD blood, including GZMK and COTL1. Focusing on tumor-specific expansion-related genes, we observed a moderate (over)expression of genes associated with dysfunctionality, such as LAG-3, CTLA4, and TIGIT (**Figure 3B**). In line with these observations, upregulated pathways in expanded tumor CD8^+^ T cells were largely related to cytokine/chemokine signaling and co-stimulation, and only moderately to TCR signaling (CD3z phosphorylation, ZAP70 translocation to immune synapse), consistent with low-level antigen recognition (**Figure 3C**).

Further characterization of the functional state of CD8^⁺^ memory T cells using gene signatures of predicted tumor reactivity, activation, cytotoxicity, and exhaustion (**Table S5)** (GO:0046635, GO:0046631, GO:0001913)^11,27^ showed a mixed profile. Tumor-infiltrating CD8^+^ T cells had higher activation and exhaustion scores than blood-derived T cells, particularly the EM subset (**Figure 3D**). Also the *predicTCR* signature^28^, designed to predict tumor reactive clones, was higher in tumor-infiltrating T cells than blood, with a higher score for a subset of patients (**Figure 3D, S4H-I**). Other signatures for tumor reactivity, however, were similarly upregulated in tumor and blood T cells compared to HD blood T cells (*Harari_up & neoTCR8*; **Figure 3D**). The *harari_up* and *neoTCR8* scores were – intriguingly – even higher in blood-derived than tumor-derived expanded TEMRA CD8^+^ T cells^29,30^ (**Figure 3D**). We hypothesized that this remarkable pattern might be explained by relatively small transcriptomic differences between blood- and tumor-derived CD8^+^ T cells, possibly suggesting a lack of tumor/antigen-driven transcriptional reprogramming in neuroblastoma-infiltrating CD8^+^ T cells. To benchmark our findings against CD8^+^ T cells from a known highly immunogenic environment (melanoma), we analyzed the same gene signatures in CD8^+^ T cells in a published dataset of melanoma patient blood and tumors^31^. Indeed, this analysis revealed significantly more pronounced transcriptional reprogramming in terms of exhaustion and predicted tumor reactivity in melanoma than in neuroblastoma (**Figure 3E**). Together, this suggests that clonally expanded tumor-infiltrating CD8^+^ T cells are activated, but only a small fraction may be actively engaging tumor cells.

Tumor-infiltrating CD8^+^ T cells furthermore displayed a significantly lower cytotoxicity score in the tumor versus blood, which might indicate impaired functionality (**Figure 3D, S4H-I).** To determine whether the reduced cytotoxicity reflected a general tissue residency program or tumor-specific effects, we compared CD8^⁺^memory T cells from tumor and sample-matched tumor-adjacent healthy adrenal tissue of two patients **(Figure S4J)**. The cytotoxicity-related genes CTSW, FGFBP2, GZMH, KLRD1, and GNLY displayed consistently lower expression in tumor-infiltrating T cells compared to normal adrenal gland-infiltrating T cells, suggesting that loss of cytotoxic features is likely induced by tumor-specific, local cues (**Figure 3F**). Overall, these data suggest that clonally expanded CD8^+^ T cells in neuroblastoma are activated, yet with limited transcriptomic evidence of exhaustion/prolonged tumor reactivity, and with reduced cytotoxic capacity.

### IV. Clonally expanded ICOS^+^ CD4^+^ αβT cells display transcriptomic profiles of tumor reactivity

Given the tumor-specific expansion and anti-tumorigenic phenotypic features observed within the CD4^⁺^ memory T cell compartment - particularly among CD4^⁺^ICOS^⁺^ cells expressing high levels of CXCL13, one of the strongest biomarkers for functional tumor reactivity^32,33^ - we hypothesized that they might play a central role in anti-neuroblastoma responses.

To elucidate tumor-specific transcriptomic profiles among clonally expanded CD4^+^ memory T cells, we compared gene expression profiles between expanded and non-expanded clones, in tumor versus blood (**Figure 4A, S5A, Table S6-8**)^26^. In contrast to CD8^+^ T cells (**Figure 3B**), we observed a low correlation between transcriptomic profiles of clonally expanded CD4^+^ T cells in tumor and blood, and a high number of tumor-specific upregulated genes, suggesting a highly tumor-specific differentiation of the CD4^+^ memory compartment (**Figure 4A, Figure S5B-C**). This profile was mostly driven by the ICOS^+^ CD4^+^ T cell subset, concomitant with a pronounced Tfh profile (IL21, CCL21, CXCL13**; Figure S4A, S5A, C**). Moreover, tumor-infiltrating expanded CD4^+^ memory T cells displayed high levels of TOX, TOX2 and inhibitory receptors PDCD1, CTLA4, TIGIT, and LAG3 **(Figure 4A)**, associated with dysfunctionality and tumor reactivity^34^. In line with this finding, pathway analysis of upregulated genes in expanded versus non-expanded CD4^+^ memory T cells showed elevated TCR-mediated activation and IFN signaling, likely indicative of antigen recognition (**Figure 4B**).

**Figure 4:**
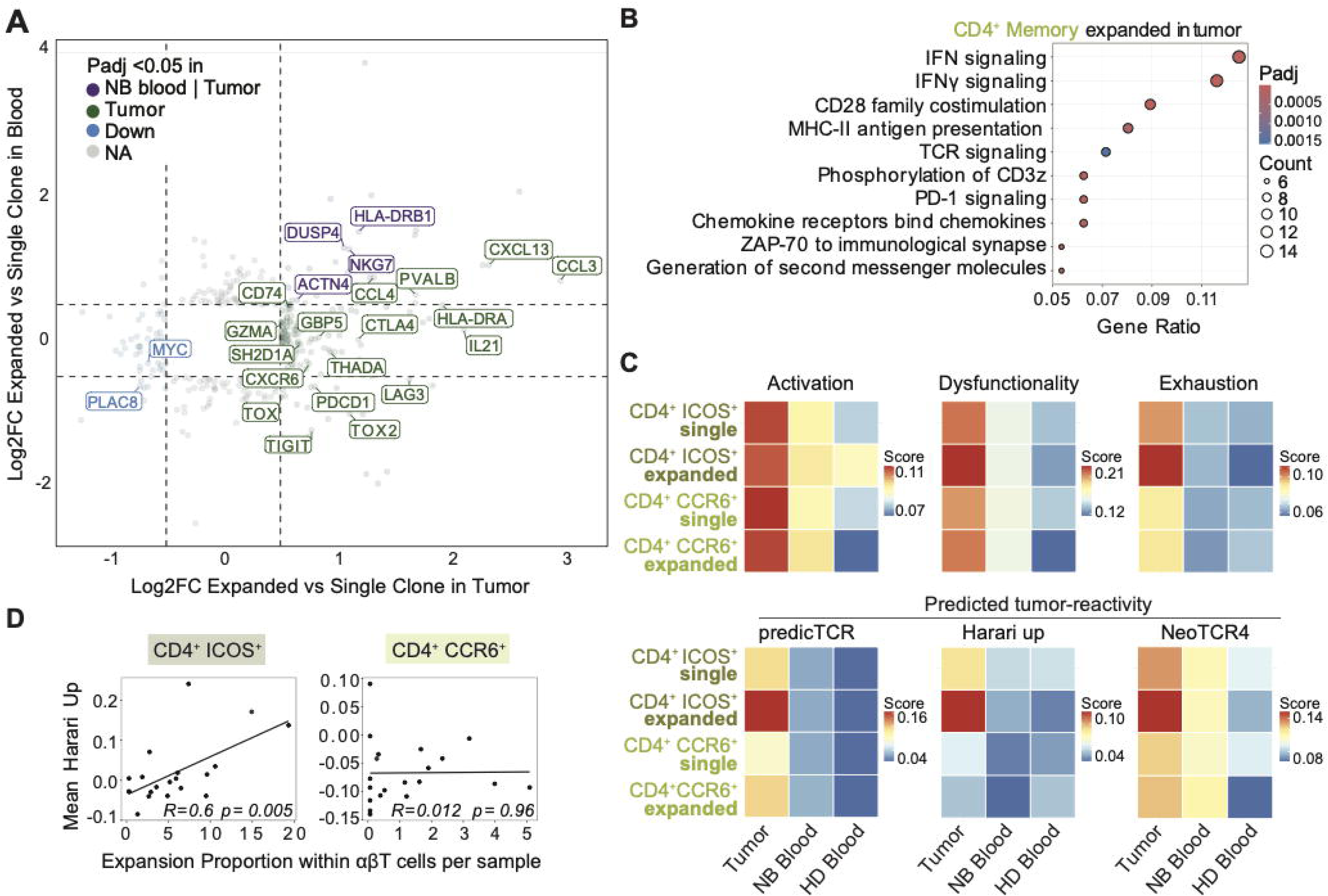
CD4^+^ memory T cells, are activated, highly dysfunctional and predicted tumor reactive. **(A)** Crater plot of significant differentially expressed genes of tumor expanded vs single clones within the CD4^+^ memory subsets. Average log2 foldchange in tumor and blood samples of these genes is shown. Green indicates genes upregulated only in expanded clones in the tumor; lilac indicates genes upregulated in expanded clones both in tumor and NB blood. **(B)** Pathway analysis showing Reactome pathways enriched in tumor expanded clones within the CD4^+^ memory. **(C)** Average value of activation, dysfunctionality, exhaustion and predicted tumor reactivity (by predicTCR, Harari_up and NeoTCR4 scores), within single and expanded CD4^+^ memory subsets in tumor, NB and HD blood. **(D)** Correlation between the tumor reactivity “Harari_up” score with the expansion proportion within αβT cells per each sample for the CD4^+^ memory subsets. Each dot indicates a sample.

To further validate potential tumor-reactivity, we evaluated established gene signatures of activation, exhaustion, dysfunction, and tumor reactivity (GO:0046631)^11,27–30,35,36^. Both ICOS^+^ and CCR6^+^ tumor-infiltrating CD4^+^ T cells displayed higher signatures of activation and dysfunctionality than blood T cells. Yet, all three signatures associated with tumor reactivity (*predicTCR, neoTCR4, harari_up*) were highest among clonally expanded ICOS^+^CD4^+^ T cells (**Figure 4C, S5D-E, Table S5**). Moreover, the *harari_up* reactivity score significantly correlated with clonal expansion, specifically in ICOS^+^CD4^+^ T cells, suggesting a strong association between prolonged target recognition and clonal expansion (**Figure 4D**). Although direct tumor-reactivity of CD4^+^ T cells has been described for MHC-II^+^ tumors, neuroblastoma cells express low levels of MHC-II molecules^37^, and it is therefore unlikely to occur in neuroblastoma patients (**Figure S5F-G**). Moreover, cytotoxic features, except for GZMA, were low in CD4^+^ T cells, rendering direct cytotoxicity by this subset unlikely **(Figure S5H)**. Rather, CD4^+^ T cells have been described to control MHC-I/II negative tumors through interplay with myeloid cells^16^. Indeed, interaction analysis showed co-stimulatory interactions between myeloid cells and CD4^+^ cells in the TME **(Figure S5I)**. Collectively, these data indicate a highly activated and Tfh-differentiated profile in CD4^+^ ICOS^+^ memory T cell compartment, with high expression of tumor reactivity signatures. These cells likely contribute to tumor-immunity through B cell stimulation in the context of immature TLS structures and through interactions with the myeloid compartment. Yet, the limited evidence of direct tumor engagement in both CD8^+^ and CD4^+^ memory T cells raises the question whether other T cell subsets may be more apt to directly engage tumor cells in neuroblastoma.

### V. Vδ1^+^ γδ T cells display gene signatures of TCR engagement

Next to αβT cells, γδT cells can serve as key cytotoxic effector cells in the TME, as several recent studies in adult MHC-I^low^ tumors have indicated^14,38,39^. To investigate their potential contribution to neuroblastoma immunity, we investigated their transcriptomic profile *in situ*.

Sub-clustering of γδTCR-bearing cells yielded 9 functional clusters (**Figure 5A, S6A-B**). Analysis of γδT cell subsets by their delta-chain, previously recognized to define functionally distinct subsets, revealed that KIR^hi^ tissue-interacting, AREG^+^ activated and naïve clusters mostly consisted of TRDV1^+^ and TRDV3^+^ cells, while TRDV2^+^ cells were predominantly represented in the Th17-like circulating and CX3CR1^+^ cytotoxic circulating clusters (**Figure 5B-C**). Based on a published γδT cell activation score (GO:0046643), TRDV1^+^ and TRDV3^+^ cells were the most activated clusters **(Figure 5D-E)**. TRDV2^+^ cells were poorly activated both in the tumor and in periphery, which was consistent across patients (**Figure 5C-E, S6C**). Since functional clusters were strongly driven by delta-chain usage (**Figure 5B**), we next investigated the frequency and functionality of the delta chain-based T cell subsets. As expected from physiological tissue tropism^38,39^, TRDV2^+^(Vγ9^+^) cells were the major γδT cell subset in the blood, and TRDV1^+^ and TRDV3^+^ cells were significantly enriched within the tumor, with diverse gamma chain expression (**Figure 5F-G**). Following chemotherapy, TRDV1^⁺^γδT cells became further enriched at the expense of TRDV2^⁺^γδT cells in both tumor tissue and peripheral blood, suggesting preferential expansion and/or tumor infiltration of TRDV1^⁺^cells, or greater chemotherapy sensitivity of TRDV2^⁺^ cells, although this has not been reported before (**Figure 5F-G**). Despite their enrichment in the tumor, the average clonal expansion rate (≥2 TCR copies) of TRDV1^+^ cells was similar to that of the other TRDV subsets (∼10%), and displayed significant variability between tumor samples (**Figure 5H-I**). Similar clonal expansion rates were observed in a publicly available bulk γδTCR sequencing data of pediatric HD blood^38^ (**Figure S6D-E**).

**Figure 5:**
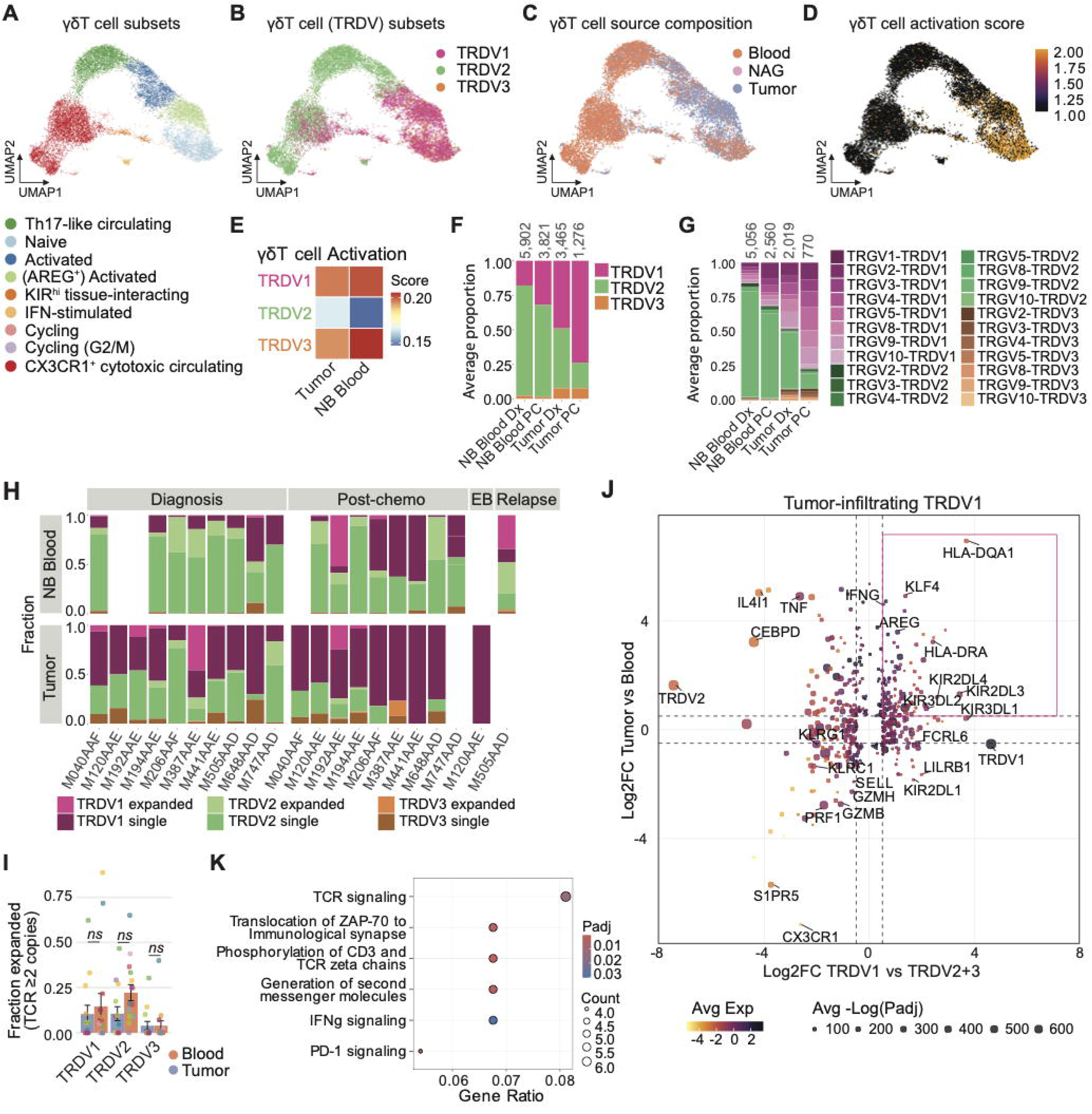
TRDV1 γδ T cells show signs of activation and TCR engagement. **(A)** UMAP dimensionality reduction of γδT cell subsets colored by annotation. **(B)** UMAP dimensionality reduction of γδT cell subsets colored by TRDV chain usage. **(C)** UMAP dimensionality reduction of γδT cell subsets colored by sample type. **(D)** gdT cell activation score overlayed to gdT cell UMAP. **(E)** Average value of activation score within TRDV subsets in tumor and NB blood. **(F)** Average proportion of TRDV1 (pink), TRDV2 (green) and TRDV3 (orange) – expressing gdT cells in NB blood and tumor samples at diagnosis and post-chemo. *Dx: diagnosis; PC: post-chemotherapy.* **(G)** Average proportion of gdT cell subsets (by gamma and delta chain composition) in NB blood and tumor samples at diagnosis and post-chemo. **(H)** Average proportion per patient, per sample type and per timepoint of expanded and single TRDV1 (pink), TRDV2 (green) and TRDV3 (orange) clones. *Dx: diagnosis; PC: post-chemotherapy.* **(I)** Average expansion proportion of TRDV cell clones within NB blood (red) and tumor (blue). **(*J*)** Crater plot of significant genes (adjusted p-value ≤ 0.05) of FindAllMarkers analysis on TRDV1 cells in the tumor (x-axis) and DGEA of tumor vs blood TRDV1 (y-axis). Color of the dot indicates average expression in the tumor. Size of the dot indicates average of adjusted p-value form the two analyses. **(*K*)** Pathway analysis showing Reactome pathways enriched in tumor TRDV1 cells.

Considering the limited expansion rate of γδT cells that could have indicated antigen-experienced cells, we evaluated their specific transcriptomic profiles within the tumor in comparison to other TRDV subsets, and in tumor versus blood, to identify key tumor-associated markers. TRDV3^+^ γδT cell numbers were too low to robustly analyze (n=272). Tumor-infiltrating TRDV2^+^ γδT cells displayed a Type 3-like program (IL17RE^+^ CCR6^+^ IL23R^+^) and were enriched for interleukin and chemokine signaling pathways, which could point to a tumor-promoting phenotype (**Figure S7A-C, Table S9-10**). Their specific expression of CXCR6 and CCR6, combined with the predicted CXCR6-CXCL16 interaction axis suggest recruitment into the tumor tissue by tumor-associated myeloid cells^40^ (**Figure S7D-F**). In contrast, tumor-derived TRDV1 cells expressed transcripts for IFNG, the inhibitory receptors KIR2DL4, KIR2DL3, KIR3DL1, KIR3DL2 and FCRL6, and the activation markers HLA-DRA and HLA-DQA1, suggestive of an activated state (**Figure 5J, S7G-H, Table S9,11**). Genes specifically upregulated in tumor-infiltrating TRDV1^+^ γδT cells were enriched for TCR signaling signatures and downstream IFN-γ/PD-1 signaling, suggesting active antigen recognition in the TME (**Figure 5K**). Together, these data suggest bystander recruitment of Vδ2^+^ γδT cells and an activated, putatively tumor reactive profile of Vδ1^+^ (and possibly Vδ3^+^) γδT cells in the neuroblastoma TME.

### VI. Functional TCR screening identifies a fraction of neuroblastoma-reactive αβ- and γδT cells

Our results so far suggest that only a small fraction of neuroblastoma-infiltrating CD8^+^ T cells may be actively engaging tumor cells (**Figure 3E, S4H**), while Vδ1^+^ γδT cells displayed more pronounced signs of tumor reactivity (**Figure 5E, J, K**). To elucidate whether a fraction of these T cell subsets in the neuroblastoma TME indeed bears neuroblastoma-specific TCRs, we selected 48 αβTCRs and 6 Vδ1^+^ γδTCRs from the tumors based on high tumor reactivity scores (αβTCRs only) and/or high clonal expansion rates, and transduced them into Jurkat reporter cells (**Figure 6A**). Functional screening of co-cultures of αβTCR-transduced Jurkats with HLA-matched neuroblastoma cell lines recorded 2 out of 48 putatively tumor reactive TCRs responding to one or more neuroblastoma cell lines, based on increased expression of activation marker CD69 (**Figure 6B,C**). Co-culture of γδTCR-transduced NFAT-GFP reporter Jurkats showed that these γδTCRs showed differential reactivity patterns to the 6 tested neuroblastoma cell lines with 2 out of 6 TCRs, both derived from the same patient, responding strongly to cell line SKNAS (**Figure 6D,E**). Together, these data show that a fraction of neuroblastoma-reactive αβTCRs and γδTCRs can be identified within the TME, which is a remarkable finding for αβTCRs considering the low MHC-I expression of neuroblastoma cells.

**Figure 6:**
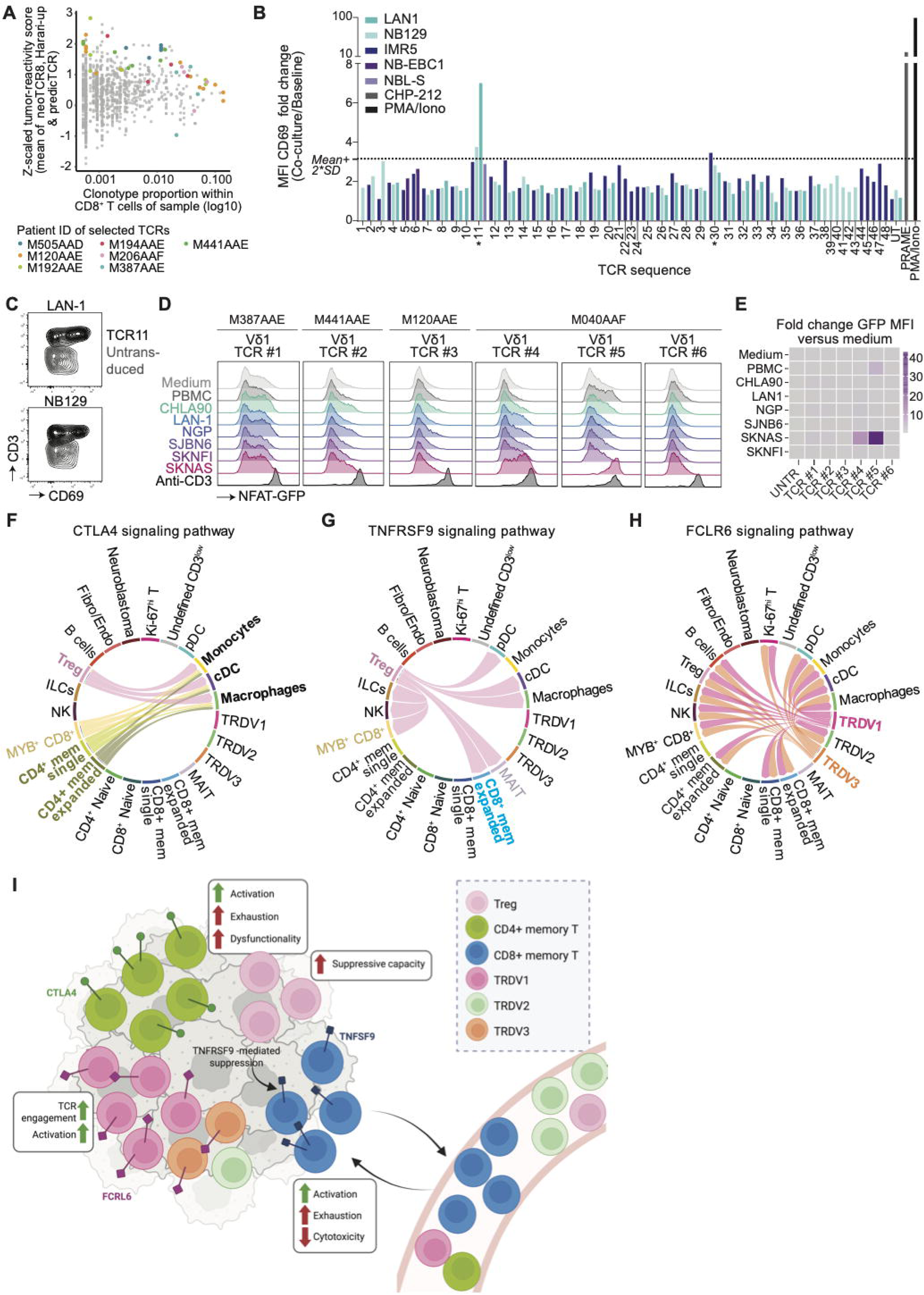
A fraction of tumor-infiltrating expanded αβ- and γδT cells is reactive to neuroblastoma cell lines *in vitro*. **(A)** Combined tumor reactivity score (mean of within-sample z-scored neoTCR8, Harari_up and PredicTCR scores) and within-sample clonotype proportion of CD8^+^ T cells of all tumor-infiltrating CD8^+^ unique T cell clones in our dataset. Colored dots indicate TCRs selected for *in vitro* screening. Color of the dot relates to individual patients. **(B)** CD69 median fluorescent intensity (MFI)-fold changes in CD8+ TCR-transduced Jurkat cells co-cultured with different neuroblastoma cell lines, normalized to the average baseline CD69 MFI of all TCR-Jurkat cells in medium only. For TCR-Jurkat cell lines #1-6 (with all target cell lines) and #11 (only with IMR5 and NBL-S co-culture) the CD69 MFI was normalized to their respective TCR-Jurkat cell line baseline CD69 MFI in medium only. A threshold for statistically significant upregulation of CD69 was set at the mean CD69 MFI-fold increase + 2x the standard deviations, for all conditions (excluding positive and negative controls), depicted by the dotted line. TCR sequences exceeding the threshold are indicated with *. **(C)** CD69 expression in TCR11-transduced vs untransduced CD8+ Jurkat cells. Gated on viable single cells. **(D)** NFAT-GFP signal in Jurkat cells transduced with selected Vδ1 TCRs, co-cultured with different neuroblastoma cell lines, medium only, healthy donor PBMCs or activated with anti-CD3. **(E)** NFAT-GFP (GFP/CD69, whichever will be chosen) MFI-fold increases in Vδ1 TCRs-transduced Jurkat cells co-cultured with different neuroblastoma cell lines, medium only, healthy donor PBMCs or activated with anti-CD3, normalized to their respective Vδ1 TCR-Jurkat cell line baseline (GFP/CD69) MFI in medium only. Reactivity of the two gdTCRs was identified based on >10-fold increase in NFAT-GFP signal compared to medium control. **(F)** CTLA4 signaling pathway network. Dimension of connecting line indicates weight of probability of the interaction. **(G)** TNFRSF9 signaling pathway network. Dimension of connecting line indicates weight of probability of the interaction. **(H)** FCRL6 signaling pathway network. Dimension of connecting line indicates weight of probability of the interaction. **(I)** Schematic representation of T cell dynamics and subset-specific immune checkpoint targets in neuroblastoma.

Having functionally confirmed the presence of tumor-reactive T cells in the neuroblastoma TME, we questioned how TME-infiltrating T cell subsets could be targeted to boost responses against neuroblastoma, since they may be restricted by inhibitory interactions with e.g. Tregs and myeloid cells. While immune checkpoint inhibition with PD-1 antibodies has so far proven unsuccessful in pediatric solid tumors^10^, targeting alternative inhibitory interactions which are more specific to the key effector subsets may be more effective. We therefore analyzed the expression of established and putative (ITIM motif–containing^41^) inhibitory receptors, together with intracellular mediators enriched in tumor samples compared to patient and healthy donor blood. The inhibitory receptor HAVCR2 (TIM-3) and CD74, the receptor for the neuroblastoma immunomodulatory factor MIF^12^, were similarly expressed across effector T cell subsets (**Figure S8A**). Other genes displayed more subset-specific expression. CTLA4, TOX2 and NDFIP2 were characteristic of CD4^+^ memory T cells, with the highest expression observed in CD4^+^ ICOS^+^ T cells (**Figure 6F, S8A,B).** For CD8^+^ T cells, the TNFSF9-TNFRSF9 (4-1BBL-4-1BB) interaction with Tregs stood out, with specific higher 4-1BBL expression in expanded CD8^+^ EM T cells (**Figure 6G, S8A, S9**). 4-1BBL triggering can directly inhibit CD8 signaling^42^. Moreover, Treg-expressed and CD8^+^-expressed 4-1BB can compete for co-stimulatory signaling by antigen-presenting cells^43^. Within γδT cells, LAG3 was expressed across all subsets (**Figure S8A, S10**). The inhibitory receptors FCRL6^44^ and several KIRs showed particularly high expression in TRDV3^+^ and expanded TRDV1^+^ cells, which we identified as putative key tumor-reactive cells^45^ (**Figure 6H, S8A, S10**).

Together, we functionally confirmed the presence of tumor-reactive TCRs in the neuroblastoma TME and defined subset-specific immune checkpoints that are highly expressed in expanded, likely tumor reactive T cell subsets, providing a model for anti-tumor immunity in neuroblastoma, and a framework for actionable strategies to enhance T cell responses (**Figure 6I**).

## Discussion

Low MHC-I levels and the lack of effective immune checkpoint therapies for neuroblastoma pose the question whether T cell responses are mounted at all, and if so, which mechanisms may impede tumor eradication^6,10^. To answer this question, we generated a comprehensive pediatric single-cell atlas of T cell dynamics in neuroblastoma. We revealed accelerated tumor-driven differentiation of circulating αβT cells; mapped connections between clonal dynamics and transcriptomic profiles of αβ and γδT cells during treatment across tissues; and delineated tumor reactive αβ and γδT cell subsets and their specific targetable immunosuppressive axes.

While previous studies reported little clonal expansion of T cells in neuroblastoma patients^46^, here we reveal a substantial fraction of clonally expanded memory T cells, already at diagnosis. By relating clonal expansion to functional properties, we show that most likely, Vδ1^+^ γδT lymphocytes are the main direct tumor-engaging T cell subset in neuroblastoma, while only a small fraction of CD8^+^ T cells appears to directly engage tumor cells. Although CD8^+^ memory T cells are classically considered the main tumor-reactive T cell subset^47^, we observed among CD8^+^ T cells an intriguing combination of 1) accelerated peripheral naive-to-memory conversion, with 2) similar activation profiles and clonal expansion rates between blood and tumor, 3) only limited exhaustion/tumor reactivity signatures, 4) tumor-site specific decreased cytotoxicity and 5) 2 out of 48 screened tumor-reactive CD8^+^ TCRs. Together, this suggests classical activation of CD8^+^ T cells with tumor-reactive TCRs within the lymph nodes and subsequent migration towards to tumor, yet lack of antigen recognition at the tumor site due to low MHC-I expression, subset-specific inhibitory cues, or spatial exclusion from tumor cores^48^. Together, this leads to a lack of clear exhaustion/tumor reactivity signatures which are generally thought to be elicited by repetitive and prolonged antigen stimulation^49^. The limited direct tumor engagement by CD8^+^ T cells may explain the poor efficacy of anti-PD-1 therapy in neuroblastoma^10^, as similarly observed in MHC^low^ melanoma^50^. These findings again nominate the low MHC-I expression on neuroblastoma tumors as a key barrier for CD8^+^ T cell responses in neuroblastoma, and emphasize the need for translational strategies to upregulate tumoral MHC-I levels, such as HDAC inhibition^51,52^.

Previous studies have reported that CD4^+^ T cells can exert anti-tumor activities without the aid of CD8^+^ T cells, mediated either via direct cytotoxicity through MHC-II expression by tumor cells, or via indirect mechanisms^17^. Specifically, the absence of MHC-I expression in tumors has been recently shown to increase their susceptibility to CD4^+^ T cell cytotoxicity^53^, which can also be exerted via the myeloid compartment^16^ or via reprogramming of stromal cells towards a pro-immunogenic phenotype^54^. Considering the low MHC-II expression levels on neuroblastoma cells and the low cytotoxic profile of CD4^+^ T cells in this dataset, we consider direct tumor cytotoxicity unlikely. Rather, ICOS^+^CD4^+^ T cells may orchestrate broader tumor immunity through interactions with B cells and myeloid cells, contributing to the formation of (immature) TLS-like structure. Such immature TLS-like structures were recently described in neuroblastoma, with a more pronounced loss of spatial structure in more aggressive neuroblastoma tumors^26,52^. This highlights that not only CD8^+^ T cell responses, but also CD4^+^ T cell responses are suboptimally formed in the more aggressive tumors.

Given their MHC-I independence^55^, TRDV1 γδT cells are known potent anti-tumoral effector cells in MHC-I^-^ tumors, like β2-microglobulin-deficient colon cancers^14^. In our dataset we observed a pronounced activation profile with signatures of TCR engagement, suggesting a role in anti-neuroblastoma responses, which we functionally validated in 2 out 6 analyzed γδTCRs. Moreover, we previously demonstrated a functional role for γδT cells with *ex vivo* expanded neuroblastoma-derived TILs, revealing that Vδ1^+^ (and Vδ3^+^) γδT cells responded to autologous tumor tissue^45^. γδT cells were recently also found to be present in neuroblastoma tumor cores at higher frequencies than CD8^+^ T cells, supporting our finding of tumor engagement by γδT cells^56^. Because of their natural tropism to the tumor bed and cytotoxic potential even against MHC-I low tumors, Vδ1^+^ γδT could be exploited as main product for tumor-infiltrating lymphocyte (TIL) therapy^45^, e.g. by selectively expanding them from the tumors and modulating their cytotoxicity. Vδ1^+^ γδT cells have been already successfully expanded from peripheral blood, showing promising results for hematological malignancies and, recently, also for solid tumors^57,58^. While we did observe gene signatures of activation in Vδ3^+^ γδT cells and have prior evidence of tumor reactivity^45^, their low abundance prevented further analyses needed to reliably assess their contribution to anti-tumor responses.

Lastly, we identified subset-specific immune checkpoints as potential targets for immunotherapy, to enhance the effector functions of CD4^+^, CD8^+^ and Vδ1^+^ T cells. CD4^+^ memory T cells specifically expressed CTLA4. Clinical translation of CTLA-4 blockade may be feasible, since both ipilimumab and tremelimumab have received FDA/EMA approval, and ipilimumab has already been evaluated in a phase I trial in pediatric solid tumors (including one neuroblastoma patient), although no overall tumor regression was observed^59^. Moreover, TIGIT and LAG-3 were broadly expressed in tumor-infiltrating CD4^+^ and CD8^+^T cells, underlining the potential of the NECTIN2-TIGIT axis previously identified by our group and others, showing pre-clinical success for neuroblastoma tumors^11,60^. LAG-3-blocking antibody relatlimab has received FDA/EMA approval, and others are currently in phase III clinical trials^59,61^. For CD8^+^T cell-specific checkpoints, we identified the suppressive 4-1BB/4-1BBL axis between Tregs and CD8^+^ memory T cells. Selective depletion of tumor-infiltrating Tregs with anti-4-1BB/anti-CCR8 bispecific monoclonal antibodies has already shown pre-clinical success^62^. Lastly, for Vδ1^+^ (and Vδ3^+^) γδT cells, their anti-tumor activity could be enhanced with γδT cell-specific immune checkpoint blockade with anti-FCRL6. FCRL6 was previously associated with diminished lymphocyte cytotoxicity in breast cancer, and its blockade could enhance pro-inflammatory cytokine secretion *in vitro*^44^. Given the high expression of inhibitory KIR receptors on Vδ1^+^ γδ T cells, targeting these receptors may represent a promising strategy to enhance their antitumor activity. Although anti-KIR monoclonal antibodies are currently being evaluated in clinical trials for adult cancers, their therapeutic effects have primarily been investigated in the context of NK-cell activation, whereas their impact on γδ T-cell function remains largely unexplored^63,64^. Thus, while subset-specific immune checkpoint inhibition represents has a strong rationale, particularly therapeutics targeting γδ T cell inhibitory receptors are still in (pre)clinical development.

In summary, we generated a comprehensive single-cell T cell atlas of neuroblastoma tumors, which offers an in-depth characterization of the T cell landscape and uncovered a model of T cell reactivity. In addition, our findings provide novel starting points for translational strategies to enhance TIL responses in neuroblastoma. These opportunities will require functional validation to evaluate their translational potential. Moreover, ancillary studies, including spatial characterization, lacking in the current model, could help dissect the spatial interaction between T cells and other cells within the microenvironment to leverage the most relevant therapeutic targets. Combined, these insights can supply subset-specific targetable leads for the development of more potent immunotherapies for patients with neuroblastoma, and potentially, other pediatric and adult solid tumors.

## Methods

### Human samples

Included samples were obtained from 10 patients treated in the Princess Máxima Center in Utrecht from October 2021 to January 2024. Patient matched tumor and blood samples were obtained at diagnosis pre-treatment (n=10 tumor; n=8 blood), during chemotherapy (n=1 tumor), at surgical resection post-chemotherapy (n=8 tumor; n=8 blood; n = 1 bone marrow) or at tumor relapse (n=1 tumor; n=1 blood). Of these, 9 patients had matched samples pre- and post-chemotherapy. Bone marrow sample was integrated together with blood samples in the analysis, given its similarity in cluster composition and gene expression. Demographic and clinical information, including tumor staging according to INRG, INRGSS and INSS criteria, was collected (**Table S13**).

This study received approval by ethics review boards as per national legislation. Patient tumor and blood samples were obtained in accordance with institutionally approved research studies by the institutional review board of the Erasmus Medical Center Rotterdam (MEC-2016-793) and the biobank committee of the Princess Máxima Center (PMCLAB2020.093/2023.0385). The studies were conducted in accordance with the Declaration of Helsinki. Age-appropriate written informed consent from patients and/or parents was obtained prior to inclusion of each patient in the study.

Age-matched healthy donor peripheral blood mononuclear cell (PBMC) data were obtained via the dataset EGAD00001007718, with access granted by the Sanger Institute^18^, or from bulk γδ TCR repertoire data from healthy pediatric donors from the study PMC11425769, downloaded from NCBI GEO under the accession code GSE263248^38^.

### Isolation of cells from tumor samples

Tumor samples were collected in the operating room and processed within 5 hours after surgery. Material was cut into pieces of ∼1-2mm^3^ and frozen in DMEM (Gibco #22320) with 10% DMSO. For single-cell RNA sequencing, samples were thawed, minced in pieces < 1mm^3^ and enzymatically digested with 2.5 mg/mL of collagenase IV (Gibco #17104-019) + RNaseOut 1:1000 (Invitrogen #10777-019) in RPMI (Gibco #61870) with 5% fetal calf serum (FCS) at 37°C with agitation for maximum 90 minutes. Digested samples were then filtered through a 70μm cell strainer and, when necessary, suspended into single cells with NeuroCult dissociation kit (StemCell Technologies #05707), according to the manufacturer’s protocol, and washed. To enable single-cell FACS sorting of live, nucleated (DAPI-DRAQ5^+^) cells on a Sony SH800S sorter, diagnosis and relapse biopsies were only stained with DAPI (Miltenyi #130-111-570) and DRAQ5 (BioLegend #424101), as it does not require washing and thereby preserves as many as possible cells of these small samples. The larger post-treatment resections and healthy adrenal gland samples were stained with Live/Dead eFluor 506 fixable viability dye (Invitrogen #65-0866-14) followed by anti-CD45 BV421 (HI30 clone; BioLegend) and anti-CD3 PE (OKT3 clone; BioLegend) antibodies for T cell enrichment and DRAQ5 for debris exclusion. Both the CD45+CD3^+^ T cells and CD3^-^ fractions were sorted and used for the library preparation.

### Isolation of cells from blood samples

Neuroblastoma patient PBMCs were freshly isolated by Ficoll-Paque (Cytiva #17144003) density centrifugation and frozen in FCS with 10% DMSO. For single-cell RNA seq, samples were thawed and stained as the previously described post-treatment resection samples. Live CD45^+^CD3^+^ and CD3^-^ fractions were FACS sorted on a Sony SH800S sorter and used for the library preparation.

### Single-cell library construction and sequencing

For diagnosis and relapse biopsies, a maximum of 25,000 sorted cells was resuspended in PBS 0.04% BSA at a density of 700-1200 cells/μL and used for library preparation. Tumor resection and blood samples were enriched for T cells by pooling together a maximum of 20,000 CD3^+^ T cells with a maximum of 5,000 CD3^-^ cells.

Libraries for single-cell RNA and αβTCR sequencing were generated using the Chromium Single Cell 5′ v2 dual-index Gel Bead and Library Construction kit, and the Chromium Single Cell Human TCR Amplification Kit (10x Genomics, CA, USA, # PN-1000190 and # PN-1000252, protocol number CG000331) according to the user guidelines. For γδTCR sequencing, 2μL of amplified cDNA was used to amplify γδTCR sequences making use of previously published custom primers^65^. Briefly, the first amplification step was performed with the forward primer: GATCTACACTCTTTCCCTACACGACGC and the outer reverse primers TRGC: ATCCCAGAATCGTGTTGCTC and TRDC: CCCACTGGGAGAGATGACAA, while for the second target enrichment the same forward primer was used with the inner reverse primers TRGC: GGGGAAACATCTGCATCAAG and TRDC: GACAAAAACGGATGGTTTGG. Agilent Bioanalyzer High Sensitivity DNA chips were used to check for quality of the libraries and DNA concentration using a Bioanalyzer 2100 machine (Agilent Technologies). DNA concentration was also measured using Qubit 1x dsDNA high sensitivity kit (ThermoFisher #Q33230). Indexed libraries were pooled and sequenced on Illumina NovaSeq 6000 (GEX and αβTCR) or NovaSeq X Plus (γδTCR) device.

### Single-cell data alignment

For both the patient and the age-matched healthy donor data, the sequencing data of the pooled samples was aligned to the GRCh38 genome using CellRanger software from 10x genomics (v7.2.0) in the *cellranger multi* mode. The GRCh38 reference genome and GRCh38 V(D)J reference provided by CellRanger were used for the scRNA and αβ/γδTCR data, respectively. For the healthy donor data, Souporcell (v2.5) was used to demultiplex the pooled sample data in reference-free mode. Subsequently, the reference vcf file was used to assign the souporcell clusters to the selected donors (NP26, NP18, NP23, NP28, NP39, NP31, NP13, NP16, NP37, NP19).

### Single-cell data processing and quality control

Count matrices were loaded into Rstudio (v4.4.0), which was used for all downstream analyses unless otherwise mentioned. Doublets were identified using scDblFinder (v.1.16.0) and removed. Low-quality cells were excluded using a cutoff of nCount ≤ 1000 and nFeature ≤ 25. Dead cells were filtered out by evaluating the percentage of expressed mitochondrial genes ≥ 20%. Downstream analyses were implemented using Seurat (v.5). Data was normalized by Seurat’s log normalization. Cell cycle phase, mitochondrial, sex-related and stress-related genes as well as TCR and B-cell receptor (BCR) genes were removed from the highly variable features to avoid clustering based on these cell-specific processes. Principal components (PCs) were calculated using ‘RunPCA’ and by using ‘ElbowPlot’ visualization, the first 30 principal components were input for ‘RunUMAP’ function to generate a UMAP bidimensional representation. To compute neighboring clusters, FindNeighbors and FindClusters were used with 30 dimensions and a resolution of 1.5, respectively. Lastly, data was batch corrected using the RunHarmony function of the harmony (v.1.2.0) package. For γδTCR HD sequencing data of donors HDL124, COV060, AUD181, AUD190, HDL111, AUD194, AUD207, HDL072, AUD182, AUD192, AUD205 and COV053, data was filtered to contain only in-frame δ-chain V(D)J rearrangements with no premature stop codons. Repertoires were down sampled to match the NB blood sample size, with clone calls based on the amino acid sequence of the δ CDR3. For the melanoma dataset, the publicly available “HumanIntegratedBlood” and “HumanIntegratedTumor” datasets were downloaded from GEO (accession number GSE159251). The blood and tumor datasets were subsequently normalized and integrated using the same workflow applied to the NB dataset.

### Cluster identification and annotation

To annotate the clusters, we combined evaluation of cluster-specific expression of cell lineage markers and published gene expression signatures. We used both the *projectTILs* (v3.3.0) reference dataset for tumor-infiltrating lyphocytes^66^ and the R package SingleR (version 2.4.1) in the “fine” mode to project the reference datasets *MonacoImmuneData, BlueprintEncodeData, DatabaseImmuneCellExpressionData, NovershternHematopoieticData* and *HumanPrimaryCellAtlasData*^67–71^. To further refine our annotation, we evaluated the top markers expressed per cluster with the ‘FindAllMarkers’ function using Wilcoxon-Rank sum test method and cross-referenced these with known T cell subset-specific expression profiles.

### Differential gene expression analysis

Differential gene expression analyses (DGEA) were performed to compare gene expression of cells between (1) healthy donor blood versus NB patient blood; (2) NB patient tumor versus NB patient blood; (3) normal adrenal gland versus tumor; (4) persistent clones post-chemotherapy versus at diagnosis or (5) within samples (for identification of expanded clonotype-specific markers). For these comparisons we used the “FindMarkers” function using “MAST” as test method, with the latent variables “sample_ID” or “Patient_ID”, where applicable. For identification of expanded clonotype-specific markers, to have sufficient statistical power for both expanded and non-expanded cells, CD8^+^ TEMRA, CD8^+^ EM and γδ1 T/ CD8^+^ memory cells were merged in CD8^+^ memory; CD4^+^ CCR6^+^ and CD4^+^ ICOS^+^ cells were merged in CD4^+^ memory. For crater plots, only significant differentially expressed genes (adjusted *p*-value ≤ 0.05) between expanded versus non-expanded clones in the tumor were plotted. For analysis of markers expressed by tumor-infiltrating TRDV subsets we used the “FindAllMarkers” function using “MAST” as test method, with the latent variables “sample_ID, as well as the “FindMarkers” function using “MAST” as test method, with “Patient_ID” as latent variables. Differentially expressed genes (DEGs) were selected based on an average log2-fold change (logFC) ≥ 0.5, and adjusted *p*-value ≤ 0.05 (adjusted based on Bonferroni correction using all genes in the dataset).

### Pathway analysis

For pathway analysis we made use of the Reactome pathway database^72^ using the enrichPathway() function from ReactomePA^73^. Differentially expressed genes with log2-fold change (logFC)□≥ 0.5 and adjusted *p*-value ≤ 0.05 were used as input for over-representation analysis. Upregulated pathways were defined as those with an adjusted p-value ≤ 0.05.

### Clonotype analysis

For αβ scTCR libraries, the outputs of *cellranger multi* were directly used as input for scRepertoire (v.1.12.0), where paired αβTCRs were reconstructed using ‘combineTCR’ function, removing cells without paired-chain information. αβTCR information was added to the Seurat object using the ‘combineExpression’ function. For αβ T cells we defined a clonotype based on the α-chain CDR3 nucleotide sequence paired with β-chain CDR3 nucleotide sequence, combined with the gene sequence of the full TCR chains (clonecall = “strict” in scRrepertoire). A cell is defined as expanded when ≥ 2 cells have the same clonotype.

For γδTCR libraries, the unfiltered contig annotations of *cellranger* were filtered manually, ensuring that only productive and full-length chains that were covered by at least 3 UMIs passed the filtering, as well as containing either paired γδ chains or a δ chain only. For γδ T cells, given their biologically limited TCR variability^74^ and low cell number in our dataset, we defined a clonotype based on the amino acid sequence of the δ-chain CDR3, known to be the most variable section. Moreover, despite this clonotype definition, eight samples for the γδ T cell clonotypes (and none for the αβ) had 30 or less cells and were excluded in subsequent analysis evaluating clonal proportions, as they would cause bias in the analysis (**Supplementary methods**). A cell is defined as expanded when ≥ 2 cells have the same clonotype.

### Overlap TCR repertoire to known TCR sequences

For αβT cells we evaluated the proportion of sequences with known reactivity using the VDJdb database. ^75^.

### Interaction analysis

The CellChat algorithm v2.2 was applied to perform an unbiased ligand-receptor interaction analysis. For this analysis we made use of the ChellChat curated ligand-receptor database^76^ to which we added published receptor-interaction for the following: CD81, CXCR6, TIGIT, CD74, CTLA4, LAG3, PDCD1, F2R, PTPRJ, ICOS, CD28, TNFRSF9, TNFRSF4, TNFRSF18, TNFRSF25, CD27, CD40LG, CD2 (additional receptor-interaction pairs provided in **Table S14**). Calculation of communication probability was carried out only for subsets with ≥ 35 cells, as described in the CellChat vignette using as cutoff a minimum of 10 cells for relevant interactions.

### Gene signature score analysis

CD4 activation (*CD4_act1*)^35^, CD4 dysfunctionality (*CD4_dysf1*)^27^, neoTCR CD4 and CD8 (*neoTCR4, neoTCR8*)^30^, *positive regulation of alpha beta T cell activation (GO:0046635)*, tumor reactivity (*harari*)^29^, cytotoxicity (*cytotox1*)^27^, exhaustion (*exhaustion1*)^27^ and *regulation of gamma delta T cell activation (GO:0046643)* gene lists were selected from published literature. *Alpha Beta T cell activation (GO:0046631)*, *T cell mediated cytotoxicity (GO:0001913)*, *3CA metaprogram CD4 T cell dysfunction*^36^ and *exhaustion_wienke*^11^ were selected as additional signatures to evaluate T cell activation, cytotoxicity, dysfunctionality and exhaustion. Scores were calculated using the *AddModuleScore_UCell()* function from *UCell*^77^. List of genes within each score is provided in **TableS5**.

### Tumor reactivity prediction

*PredicTCR v2*^28^ was used to annotate tumor-reactive clonotypes. Input to *PredicTCR* was the log-normalized data from the Seurat object, analyzed for each sample separately. The probability of reactivity was predicted using the *PredicTCR* model, where the threshold is determined using Jenk-Fisher natural break optimization.

### FACS data of high-risk neuroblastoma patients

Spectral flow cytometry analysis was performed on n= 10 post-chemotherapy high-risk neuroblastoma tumor samples (different than the sequenced ones) defrosted and washed. Tumor digest *ex vivo* was stained with Fixable Viability Dye eFluor 506 (65-0866-14; Thermo Fisher Scientific) for 20 min at 4°C. After washing, cells were stained with HLA-ABC-PerCP (W6/32, 311421, Biolegend) and HLA-DR-BUV661 (G46-6, 612981, BD Bioscience) antibodies in the dark for 20 min at 4°C. Samples were acquired on an Aurora flow cytometer (Cytek Biosciences) with SpectroFlo software. Cell subsets were excluded for further analysis when the cell numbers were below 20 cells. Data analysis was carried out with FlowJo Star 10.7.1.

### Culture of Cell lines

For αβTCR transduction, CD8^+^ Jurkat cells (kindly provided by Wouter Scheper, Netherlands Cancer Institute, Amsterdam) were cultured in lymphocyte medium, consisting of RPMI 1640 medium (Gibco, Cat. No. 21875-034), supplemented with heat-inactivated sterile-filtered FCS (Capricorn Scientific, Cat. No. FBS-16A), 1% P/S (Gibco, Cat. No. 15140-122) and 1% L-Glutamine (Gibco, Cat. No. 25030-024). For γδTCR transduction, a variant of the Jurkat T cell line JE6.1-NF-κB::eGFP (REF: 10.1016/j.jim.2016.01.007), not possessing the endogenous TRA and TRB chain (knock out for TRAC and TRBC as described (REF: doi: 10.1002/cti2.1216)), was kindly provided by Peter Steinberger (Vienna, Austria). This cell line was cultured in standard complete medium, consisting of RPMI (Gibco, Invitrogen), supplemented with 10% FCS (Gibco, Invitrogen), 2 mM L-glutamine (Gibco, Invitrogen), 100 IU/mL penicillin (Gibco, Invitrogen) and 100 IU/mL streptomycin (Gibco, Invitrogen). Phoenix packaging cell line was cultured in standard complete medium consisting of IMDM (Gibco, Invitrogen), supplemented with 10% FCS (Gibco, Invitrogen), 2 mM L-glutamine (Gibco, Invitrogen), 100 IU/mL penicillin (Gibco, Invitrogen) and 100 IU/mL streptomycin (Gibco, Invitrogen). HEK293T cells were cultured in DMEM supplemented with 10% FCS, 1% MEM-NEAA and 1% L-Glut without Penicillin/Streptomycin (P/S).

Neuroblastoma cell lines CHP-212 and LAN-1 were cultured in RPMI 1640 medium, supplemented with 10% FCS and 1% Pen/Strep. Neuroblastoma cell lines IMR-5, NB-EBC1, NBL-S, SKANS, NGP, SJNB6 and SKNFI were cultured in Dulbecco’s Modified Eagle Medium (DMEM; Gibco, Cat. No. 41965-039), supplemented with 10% FCS, 1% P/S, 1% Minimum Essential Medium non-essential amino acids (MEM-NEAA; Gibco, Cat. No. 1140-035) and 1% L-Glutamine. CHLA90 was cultured in Iscove’s Modified Dulbecco’s Medium (IMDM Gibco, #12440-053), supplemented with 10% FCS, 1% Pen/Strep, 1% L-Glut and 1X Insulin-Transferrin-Selenium (Gibco, #A4000046501).

Neuroblastoma tumoroid NB129 was cultured in tumoroid medium, consisting of DMEM, low glucose, GlutaMAX (Gibco, Cat. No. 21885-025), 20% Ham’s F-12 Nutrient Mix, GlutaMAX (Gibco, Cat. No. 31765-027), 2% B-27 supplement, minus vitamin A (Gibco, Cat. No. 12587-001), 1% N-2 supplement (Gibco, Cat. No. 17502-001), 1% P/S, 20 ng/mL animal-free recombinant human EGF (Gibco, Cat. No. AF-100-15), 40 ng/mL recombinant human FGF-basic (154 aa) (Gibco, Cat. No. 100-18B), 200 ng/mL recombinant human IGF-I (Gibco, Cat. No. 100-11), 10 ng/mL recombinant human PDGF-AA (Gibco, Cat. No. 100-13A), and 10 ng/mL recombinant human PDGF-BB (Gibco, Cat. No. 100-14B), as described previously^78^.

Tumoroid line NB129 was detached using Accutase (Sigma-Aldrich, Cat. No. A6964-100ML) whilst all other adherent cell lines were detached with TrypLE Express Enzyme (Gibco, Cat. No. 12605-010). Cells were incubated at 37°C, 5% CO2, and passaged twice per week, unledss stated differently.

### Generation of αβT cell receptor transduced Jurkat lines

αβT cell receptors were selected based on high clonal expansion rates and/or high tumor reactivity scores (predicTCR^28^, *Harari_up & neoTCR8*^29,30^). The latter were Z-scaled per sample. TCR constructs were designed featuring murine constant regions in a pCCL-PPT-PGK lentiviral-vector (kindly provided by the Nierkens group, Princess Máxima Centre, Utrecht) and synthesized by GenScript. A HLA-A*02 restricted, preferentially expressed antigen in melanoma (PRAME)-SLL-recognizing TCR was used as positive control TCR^79^.

To facilitate high-throughput screening, lentivirus production was optimized for 96-well plates. 1.2 × 10^4^ cells/well low-passage HEK293T cells were seeded in flat-bottom 96-well plates in DMEM supplemented with 10% FCS, 1% MEM-NEAA and 1% L-Glut without Penicillin/Streptomycin (P/S) and cultured until 60% confluency. Cells were co-transfected using FuGENE HD (Promega, Cat. No. E2311) with the αβTCR plasmids (140.9 ng/well) and helper plasmids diluted in Opti-MEM (Gibco, Cat. No. 31985070). Following 15 minutes incubation at room temperature, transfection complexes were added to the cells. 24 hours post-transfection, the DMEM medium was replaced with RPMI 1640 medium (supplemented with 10% FCS, 1% L-Glut, 1% sodium pyruvate (Gibco, Cat. No. 11360-070), and 25 mM HEPES (Gibco, Cat. No. 15630-056)). Viral supernatants were harvested at 48 and 72 hours post-transfection, pooled per TCR, and centrifuged at 1500 × *g* for 5 minutes to pellet any contaminating HEK293T cells.

CD8^+^ Jurkat cells were seeded at 10□cells/well in 96-well round-bottom plates for transduction. Plates were centrifuged at 300 × *g* for 5 minutes, and culture medium was removed. Pellets were resuspended in viral supernatants. Plates were spinoculated at 1000 × *g* for 40 minutes at room temperature and incubated overnight. Transduction efficiency was evaluated 9 days post-transduction by flow cytometric staining for CD3 (HIT3a BioLegend, #300308) and murine TCRβ (H57-597, BioLegend, #109206). All TCRs reached the minimally required transduction efficiency of 40%. Cells were passaged every 2-3 days to maintain a cell density at or below 5^10^5^ cells/mL, preventing high background CD69 expression prior to downstream screening.

### Generation of γδ TCR construct encoding plasmids

Selected paired CDR3γ and CDR3δ sequences from the single-cell TCR data were completed with the V, J and C sequences available at the IMGT website (REF: doi.org/10.1093/nar/gkab1136). The choice of TRGC1 or TRGC2 was based on the TRGJ usage in the single-cell TCR data combined with the available single-cell TRGC1/TRGC2 gene expression data. γδ TCR constructs containing the paired TRG and TRD sequences were ordered as gBlocks (IDT) and cloned into the retroviral vector LZRS using Gibson Assembly (NEBuilder HiFi DNA Assembly Master Mix; NEB). Briefly, the LZRS was linearized by digestion with BamHI (NEB) and purified using the NucleoSpin® Gel and PCR Clean-up Kit (Macherey-Nagel) according to the manufacturer’s instructions. The purified plasmid backbone was subsequently dephosphorylated using rAPid Alkaline Phosphatase (Roche). Gibson Assembly reactions were performed using the linearized vector and γδ TCR gBlock inserts. The assembled plasmids were transformed into NEB Stable High-Efficiency *Escherichia coli* competent cells (NEB) and plated on lysogeny broth (LB) agar plates supplemented with ampicillin. Following overnight incubation at 37°C, individual colonies were selected and grown in LB medium supplemented with ampicillin. Plasmid from the overnight culture was isolated using Miniprep Kit (Qiagen) and sequenced. The plasmid with the correct sequence was then used for retroviral particle production.

### Production of γδ TCR Retroviral Vectors

Viral particles were produced using the Phoenix A packaging cell line. Briefly, Phoenix A cells were seeded at 7.5 × 10^5^ cells per 6 cm dish (BD Falcon) and placed overnight in a 7% CO_2_ incubator at 37 °C. The next day, the plasmids encoding the γδ TCR constructs were transfected using calcium phosphate as follows: per 6 cm dish, 10 µg plasmid DNA was diluted in 36 µL 2 M CaCl_2_ (homemade), and subsequently nuclease-free water (Ambion) was added to a total volume of 300 µL. The DNA-CaCl_2_ solution was pipetted dropwise into a 15 mL polystyrene tube containing 300 µL 2× HEPES-buffered saline (HBS) solution (homemade) while blowing air bubbles in the 2× HBS buffer. The mixture was incubated for 15 min at room temperature and then added dropwise to the cells. Ten minutes before the addition of the DNA mixture, 2 mL medium was added to the cells containing 1 µL of a 200 mM chloroquine (Sigma-Aldrich) solution. Cells were placed in a 7% CO_2_ incubator at 37 °C overnight. Medium was refreshed on day one. At day two, the viral supernatant was harvested and was frozen at −80 °C until use.

### Generation of γδ TCR-expressing Jurkat cell line

Jurkat cells were seeded at a density of 2.5 × 10^5^ cells per well in non-tissue-culture-treated plates coated with Retronectin (TaKaRa). Viral supernatant was added, and cells were subjected to spinoculation by centrifugation at 1000g for 90 min at 32°C. Following centrifugation, cells were incubated in a 7% CO_2_ incubator at 37 °C for 48 h. Transduced cells were subsequently washed with PBS and maintained in complete RPMI medium. Transduction efficiency was evaluated by flow cytometry as the percentage of cells expressing both CD3 (clone UCHT1; BD Biosciences) and TCRγδ (clone 11F2; Miltenyi Biotec). Transduced cells were then enriched based on CD3 expression using a FACSAria III cell sorter (BD Biosciences).

### Functional screening of αβT cell receptors

For the TCR screening assays, neuroblastoma target cells were pre-stimulated with 20 ng/mL recombinant IFN-γ (BioLegend, Cat. No. 570202) for 24 hours to induce MHC-I expression. 3.75 × 10^5^ target cells were seeded into 96-well flat-bottom plates for screening at an 1:5 effector:target (E:T) ratio. Subsequently, 7.5 × 10^4^ effector cells (TCR-transduced or untransduced Jurkat cells) were added to the respective wells in a final volume of 200 μL complete RPMI medium. PRAME-TCR-transduced Jurkat cells and Jurkat cells stimulated with phorbol 12-myristate 13-acetate (PMA) (20 ng/mL; Sigma-Aldrich, Cat. No. P8139-1MG) and Ionomycin (1 μg/mL; Sigma-Aldrich, Cat. No. I0634-1MG) served as positive controls. All co-cultures were incubated for 18 hours, to allow sufficient time for antigen recognition but to prevent activation through alloreactivity. T cell activation was measured via CD69 expression through flow cytometry. Cells were washed with DPBS and stained with Fixable Viability Dye eFluor506 (1:300; Invitrogen, Cat. No. 65-0866-14) for 20 minutes at 4°C in the dark. Cells were then washed, resuspended in FACS buffer (DPBS supplemented with 2% FCS and 2 mM EDTA) and stained for 20 minutes at 4°C in the dark with antibodies for CD8 (SK1, BioLegend, #344708), CD3 (SK7 BioLegend, #344818), murine TCRβ (H57-597, BioLegend, #109206) and CD69 (FN50, Invitrogen, #407-0699-42). Data were acquired on CytoFLEX LX or S flow cytometers (Beckman Coulter). Data were analyzed in Flowjo v10 software (LLC). The median fluorescence intensity (MFI) for CD69 was used for further analysis. The threshold for reactivity was set at the mean MFI for CD69 + 2x the standard deviations, for all measured conditions combined (excluding positive and negative controls).

### Functional screening of γδT cell receptors

γδT cell receptors were selected based on the highest clonal expansion rates within a sample. For the γδTCR screening assays, 3.75 × 10^5^ target cells were seeded into 96-well flat-bottom plates for screening at an 1:5 effector:target (E:T) ratio. Subsequently, 7.5 × 10^4^ effector cells (TCR-transduced or untransduced Jurkat cells) were added to the respective wells in a final volume of 200 μL complete RPMI medium. Jurkats stimulated with plate-coated 5 µg/mL anti-human CD3 (OKT3) served as positive controls. All co-cultures were incubated for 18 hours, to allow sufficient time for antigen recognition but to prevent activation through alloreactivity. T cell activation was measured via NFAT-GFP expression through flow cytometry. Cells were washed with DPBS and stained with Fixable Viability Dye eFluor506 (1:300; Invitrogen, Cat. No. 65-0866-14) for 20 minutes at 4°C in the dark. Cells were then washed, resuspended in FACS buffer (DPBS supplemented with 2% FCS and 2 mM EDTA) and stained for 20 minutes at 4°C in the dark with antibodies for CD3 (HIT3a BioLegend, #300308), γδTCR (Miltenyi, #130-114-027) and CD69. (FN50, Invitrogen, #407-0699-42). Data were acquired on CytoFLEX LX or S flow cytometers (Beckman Coulter). Data were analyzed in Flowjo v10 software (LLC). The median fluorescence intensity (MFI) for NFAT-GFP was used for further analysis.

### Statistical analysis

Statistical tests were performed in R, using *stats V4.3.3*, *FSA V0.9.5* and ggpubr *V0.6.0*. Statistical analyses were performed using average values per sample_ID as independent observations. For comparisons between two groups, we used the Wilcoxon rank-sum test, while comparisons among three groups were conducted using the Kruskal–Wallis’s test followed by Dunn’s test. P-values were adjusted for multiple testing using the Benjamini–Hochberg (BH) procedure. Statistical significance was evaluated for p-value adjusted ≤ 0.05, unless specified differently in the figure legends.

---

**Figure S1: (A)** UMAP dimensionality reduction of all cells of patients and HD colored by annotation and by patient/HD. **(B)** Expression of the top 5 genes per each cluster of figure S1A. Color represents average expression. Size of the dot represents percentage of cells expressing the marker. **(C)** Frequency of cell subsets within patients/HD samples. Numbers on top of the bar relate to number of total cells per cluster. **(D)** Frequency HD/patient cells per each cell subset. Numbers on top of the bar relate to number of total cells per cluster.

**Figure S2: (A)** UMAP dimensionality reduction of NB blood and HD blood T cells from tumor (blue), NB blood (red) and HD blood (green) samples colored by annotation and TCR expression. **(B)** Composition of T cell subsets by NB blood (red), HD blood (green), normal adrenal gland, normal adrenal gland (pink) and tumor (blue). Numbers on top of the bar indicate number of cells. **(C)** T cell subset composition per patient/HD, per sample type (tumor, blood or normal adrenal gland) and per timepoint. EB: excision biopsy; Rel: relapse; Post-ch: post-chemotherapy; NAG: normal adrenal gland.

**Figure S3: (A)** Percentage of αβTCR^+^ and γδTCR^+^ within HD blood, NB blood, tumor. Numbers on top of the bar indicate number of cells within each sample type. **(B)** Frequency of expanded αβTCR^+^ which TCR sequence was found in the VDJdb dataset for known-specificity TCR. **(C)** Volcano plot of differentially expressed genes between persistent clones post-chemo versus diagnosis. **(D)** Reactome pathways enriched in persisting clones after chemo vs at diagnosis. **(E)** Proportion of top 10* αβT cell clonotypes in the repertoire normalized to 100% in blood and tumor at different therapy timepoints. Numbers on top of the bar indicate total number of αβT cell clones within sample. BD: blood diagnosis; TD: tumor diagnosis; BPC: blood post-chemotherapy; TPC: tumor post-chemotherapy; BRe: blood relapse; Teb: tumor excision biopsy. **(F)** Clonal overlap between different samples measured by weighted Jaccard index. **(G)** αβTCR data overlayed to gene expression UMAP of tumor, NB blood and HD blood cells. Expanded clones are indicated in purple, single clones in yellow and no αβTCR found in grey. (**H)** Occupied repertoire space by the expanded αβT cell clones in HD, NB blood, normal adrenal gland (NAG) and tumor. Numbers on top of bars indicate number of expanded cells within that sample. Samples with low cell numbers (≤ 20) are depicted on the right.

**Figure S4: (A)** Expression of TLS-associated genes in B cells, dendritic cells (DCs) and CD4^+^ ICOS^+^ cells within the tumor. **(B)** CD40 signaling pathway network. Dimension of connecting line indicates weight of probability of the interaction. **(C)** Expression of MHC-I genes in tumor and monocyte/macrophages cells within the tumor microenvironment. Color indicates level of expression. Size indicates percentage of expressing cells. **(D)** Percentage of HLA-ABC^+^ single viable myeloid and tumor cells from FACS data of n=10 neuroblastoma patients. **(E)** Heatmap of top 30 DEG in expanded vs single CD8^+^ memory cells in the tumor. **(F)** Upset plot of DEG upregulated by expanded CD8^+^ memory only in the tumor (green), in both tumor and NB blood (lilac). Numbers on top of bar indicate number of genes within each comparison. **(G)** Expression of top20 patient-specific genes (green and lilac from figure 3B within tumor-infiltrating expanded and single CD8^+^ memory. **(H)** UCell tumor reactivity, activation, dysfunctionality and exhaustion scores per sample in tumor, NB blood and HD blood for both for single/expanded CD8^+^ memory subsets. Only statistically significant comparisons are indicated. **(I)** Average value of activation, dysfunctionality and exhaustion with different scores than in figure 3D within single and expanded CD8^+^ memory subsets in tumor, NB and HD blood. **(J)** Volcano plot of differentially expressed genes for all T cells of n=2 patients with matched normal adrenal gland (pink) versus tumor (blue) samples.

**Figure S5: (A)** Heatmap of top 30 DEG in expanded vs single CD4^+^ memory cells in the tumor. **(B)** Upset plot of DEG upregulated by expanded CD4^+^ memory only in the tumor (green), in both tumor and NB blood (lilac). Numbers on top of bar indicate number of genes within each comparison. **(C)** Expression of top20 patient-specific genes (green and lilac from figure 4A) within tumor-infiltrating expanded and single CD4^+^ memory. **(D)** UCell tumor reactivity, activation, dysfunctionality and exhaustion scores per sample in tumor, NB blood and HD blood for both for single/expanded CD4^+^ memory subsets. Only statistically significant comparisons are indicated. **(E)** Average value of activation, dysfunctionality and exhaustion with different scores than in figure 4C within single and expanded CD4^+^ memory subsets in tumor, NB and HD blood. **(F)** Expression of MHC-II genes in tumor and monocyte/macrophages cells within the tumor microenvironment. Color indicates level of expression. Size indicates percentage of expressing cells. **(G)** Percentage and median fluorescent intensity (MFI) of HLA-DR^+^ single viable myeloid and tumor cells from FACS data of n=10 neuroblastoma patients. **(H)** Expression of cytotoxicity-related genes in CD8^+^ and CD4^+^ memory subsets in tumor and blood. Color indicates level of expression. Size indicates percentage of expressing cells. **(I)** Cellchat interactions between myeloid cells with CD4^+^ T cell subsets.

**Figure S6: (A)** Expression of the top 10 genes per each cluster of figure 5A. Color represents average expression. Size of the dot represents percentage of cells expressing the marker. **(B)** TRDV subset composition of clusters for figure 5A. Numbers on top of the bar indicate number if cells within each cluster. **(C)** Activation score per patient within TRDV subsets in tumor and NB blood. **(D)** Clone size proportion of single (grey) and expanded (colored) γδT cell clones within each HD blood from bulk γδTCRseq data (green), NB blood (red) and tumor (blue) sample. **(E)** Fraction of TRDV1 (pink), TRDV2 (green) and TRDV3 (orange) – expressing γδT cells in HD blood from published bulk γδTCRseq dataset and divided by each HD and in total.

**Figure S7: (A)** Crater plot of significant genes (adjusted p-value ≤ 0.05) of FindAllMarkers analysis on TRDV2 cells in the tumor (x-axis) and DGEA of tumor vs blood TRDV2 (y-axis). Color of the dot indicates average expression in the tumor. Size of the dot indicates average of adjusted p-value form the two analyses. **(B)** Expression of Th17-related genes TRDV subsets in tumor. Color indicates level of expression. Size indicates percentage of expressing cells. **(C)** Pathway analysis showing Reactome pathways enriched in tumor TRDV2 cells. **(D)** CXCR6 signaling pathway network. Dimension of connecting line indicates weight of probability of the interaction. **(E)** CXCR6 expression within the different tumor-infiltrating γδT cell subset. Each dot indicates a cell. **(F)** CXCL16 expression in tumor-infiltrating (blue) and NB blood circulating (red) monocytes. **(G)** IFNG expression in tumor-infiltrating TRDV subsets. **(H)** HLA-DRA and HLA-DRB1 expression in tumor-infiltrating TRDV subsets.

**Figure S8: (A)** Expression of known and putative immune checkpoint markers of expanded and non-expanded CD4^+^, CD8^+^ and TRDV subsets in tumor, blood and HD blood. Color indicates level of expression. Size indicates percentage of expressing cells. **(B)** CellChat receptor-ligand analysis of significant interaction predicted between CD4^+^ T cells and the TME. Color indicates communication probability.

**Figure S9: (A)** TNFRSF9 expression within tumor-infiltrating (blue), NB blood (red) and HD blood (green) Treg. Each dot represents a cell. **(B)** CellChat receptor-ligand analysis of significant interaction predicted between CD8^+^ T cells and the TME. Color indicates communication probability.

**Figure S10:** CellChat receptor-ligand analysis of significant interaction predicted between γδT cells and the TME. Color indicates communication probability.

## Figure supplementary methods

αβ and γδT cell clonal size distribution across all samples within HD (green), NB blood (red), normal adrenal gland (NAG) (pink) and tumor (blue). Dotted line indicates threshold of 30 cells per sample.

## Supplementary Tables

All Supplementary tables have been deposited in Zenodo and are publicly available at https://doi.org/10.5281/zenodo.24160343 as of the date of publication.

**Table S1:** “FindAllMarkers” of αβTCR^+^ and γδTCR^+^ T cell clusters in tumor samples

**Table S2:** Differential gene expression analysis (“MAST” as test method, with the latent variables “sample_ID”) expanded versus non-expanded CD8^+^ memory T cells (CD8^+^ memory, CD8^+^ EM and CD8^+^ TEMRA clusters) in tumor.

**Table S3:** Differential gene expression analysis (“MAST” as test method, with the latent variables “sample_ID”) expanded versus non-expanded CD8^+^ memory T cells (CD8^+^ memory, CD8^+^ EM and CD8^+^ TEMRA clusters) in neuroblastoma blood.

**Table S4:** Differential gene expression analysis (“MAST” as test method, with the latent variables “sample_ID”) expanded versus non-expanded CD8^+^ memory T cells (CD8^+^ memory, CD8^+^ EM and CD8^+^ TEMRA clusters) in healthy donor blood.

**Table S5:** List of genes used for Ucell scores throughout the paper.

**Table S6:** Differential gene expression analysis (“MAST” as test method, with the latent variables “sample_ID”) expanded versus non-expanded CD4^+^ memory T cells (CD4^+^ ICOS^+^ and CD4^+^ CCR6^+^) in tumor.

**Table S7:** Differential gene expression analysis (“MAST” as test method, with the latent variables “sample_ID”) expanded versus non-expanded CD4^+^ memory T cells (CD4^+^ ICOS^+^ and CD4^+^ CCR6^+^) in nb blood.

**Table S8:** Differential gene expression analysis (“MAST” as test method, with the latent variables “sample_ID”) expanded versus non-expanded CD4^+^ memory T cells (CD4^+^ ICOS^+^ and CD4^+^ CCR6^+^) in healthy donor blood.

**Table S9:** “FindAllMarkers” (“MAST” as test method, with the latent variables “sample_ID”) of TRDV1^+^, TRDV2^+^ and TRDV3^+^ γδT cells in tumor samples.

**Table S10:** Differential gene expression analysis (“MAST” as test method, with the latent variables “patient_ID”) TRDV2^+^ γδT cells tumor versus blood.

**Table S11:** Differential gene expression analysis (“MAST” as test method, with the latent variables “patient_ID”) TRDV1^+^ γδT cells tumor versus blood.

**Table S12:** Differential gene expression analysis (“MAST” as test method, with the latent variables “sample_ID”) expanded versus non-expanded Tregs.

**Table S13:** Patient characteristics.

**Table S14:** Complete list of receptor-ligand interactions in the CellChat database.

## Resource availability

### Lead contact

Further information and requests for resources and reagents should be directed to and will be fulfilled by the lead contact, Judith Wienke.

### Materials availability

This study did not generate new unique reagents.

### Data Availability Statement

Single-cell RNA and TCR-seq data have been deposited at EGA as EGAS00001008549 and are publicly available as of the date of publication. This paper analyzes existing, publicly available data, accessible at EGAD00001007718 (pediatric healthy donor single-cell αβTCR data) with access granted by the Sanger Institute; at GEO under the accession code <u>GSE263248</u> (healthy pediatric donor bulk γδ TCR repertoire data); and at GEO under the accession code GSE159251 (melanoma dataset).

The Seurat objects reported in this study cannot be deposited in a public repository because of ongoing patent-related proceedings. To request access, contact the lead contact.

All original code has been deposited at Zenodo and is publicly available at DOI:10.5281/zenodo.24160018 as of the date of publication. Any additional information required to reanalyze the data reported in this paper is available from the lead contact upon request.

## Acknowledgments

We would like to acknowledge our FACS facility, particularly Niels Groenen and Jeroen van Velzen, and the Single Cell Genomics Facility, especially Aleksandra K. Balwierz, Wim J. de Jonge, Jeff DeMartino, Philip Lijnzaad and Thanasis Margaritis. We thank the Genome diagnostics section of the department of Genetics, UMC Utrecht for additional sequencing services. We thank Peter Steinberger (Medical University of Vienne, Austria) for providing a TCR-deficient version of the Jurkat T cell line JE6.1-NF-κB::eGFP.

This work received funding from the European Union’s Horizon 2020 research and innovation program under Marie Skłodowska-Curie grant agreement, No. 956285 (VAGABOND), Villa Joep, and Kika (grant number 491). D.V. received funding from WELBIO investigator program (WELBIO-CR-2022A–15, to D. Vermijlen) and a PDR-WEAVE grant (FNRS, T.0322.25 PDR FRSM, to D. Vermijlen). S.D.M is supported by the FWO postdoctoral fellowship junior 12AP724N. B.V. is supported by the FWO Research project G0A1D24N. T.L.C is supported by the HECTOR Kick-Start Seed Funding “C-3PO”. The Single Cell Genomics Facility is supported by the Children Cancer Free Foundation (KiKa).

## Author contributions

Conceptualization: E.Z., J.W. Data curation: E.Z., N.H.V, S.M., A.L.B., M.C.W., E.W.G., D.V., J.J.M, J.W. Formal analysis: E.Z., N.H.V, S.M., G.S.S., A.L.B., J.W., Funding acquisition: E.Z., S.D.M., B.V., C.L.T., A.K.B., W.J.J., P.L., T.M., M.C.W., E.W.G., D.V., J.J.M., J.W., Investigation: E.Z., N.H.V., S.M., F.P.M., S.A.A., S.D.M., B.V., C.L.T., A.L.B. Methodology: E.Z., N.H.V., S.M., F.P.M., G.S.S., S.A.A., S.D.M., B.V., C.L.T., A.K.B., W.J.J., P.L., T.M., A.L.B., A. FW.S., R.K., M.P.D., G.AM.T., E.W.G., W.S., D.V., J.W. Administration: E.Z., N.H.V., A. FW.S., R.K., M.P.D., G.AM.T., J.J.M, J.W. Resources: E.Z., N.H.V., F.P.M., G.S.S., S.A.A., S.D.M., B.V., C.L.T., A.K.B., W.J.J., P.L., T.M., A.L.B., A. FW.S., R.K., M.P.D., G.AM.T., E.W.G., M.C.W., W.S., D.V., J.J.M, J.W. Software: E.Z., N.H.V., G.S.S, C.L.T., P.L., E.W.G., J.W. Supervision: E.Z., T.M., M.C.W., E.W.G., D.V., J.J.M., J.W. Validation: E.Z., S.M. Visualization; E.Z., N.H.V., S.M., A.L.B., J.W. Writing – draft: E.Z. Writing – review and editing: N.H.V., S.M., M.C.W., E.W.G., J.J.M, J.W. All authors critically revised the manuscript.

## Declaration of generative AI and AI-assisted technologies in the manuscript preparation process

During the preparation of this work, the author(s) used ChatGPT in order to assist with language editing, including sentence rephrasing, improving clarity and readability. After using this tool/service, the author(s) reviewed and edited the content as needed and take(s) full responsibility for the content of the published article.

