## Supplementary figures and images for "Single-cell T cell receptor sequencing deciphers αβ- and γδ-T cell responses in neuroblastoma"

### Figure S1

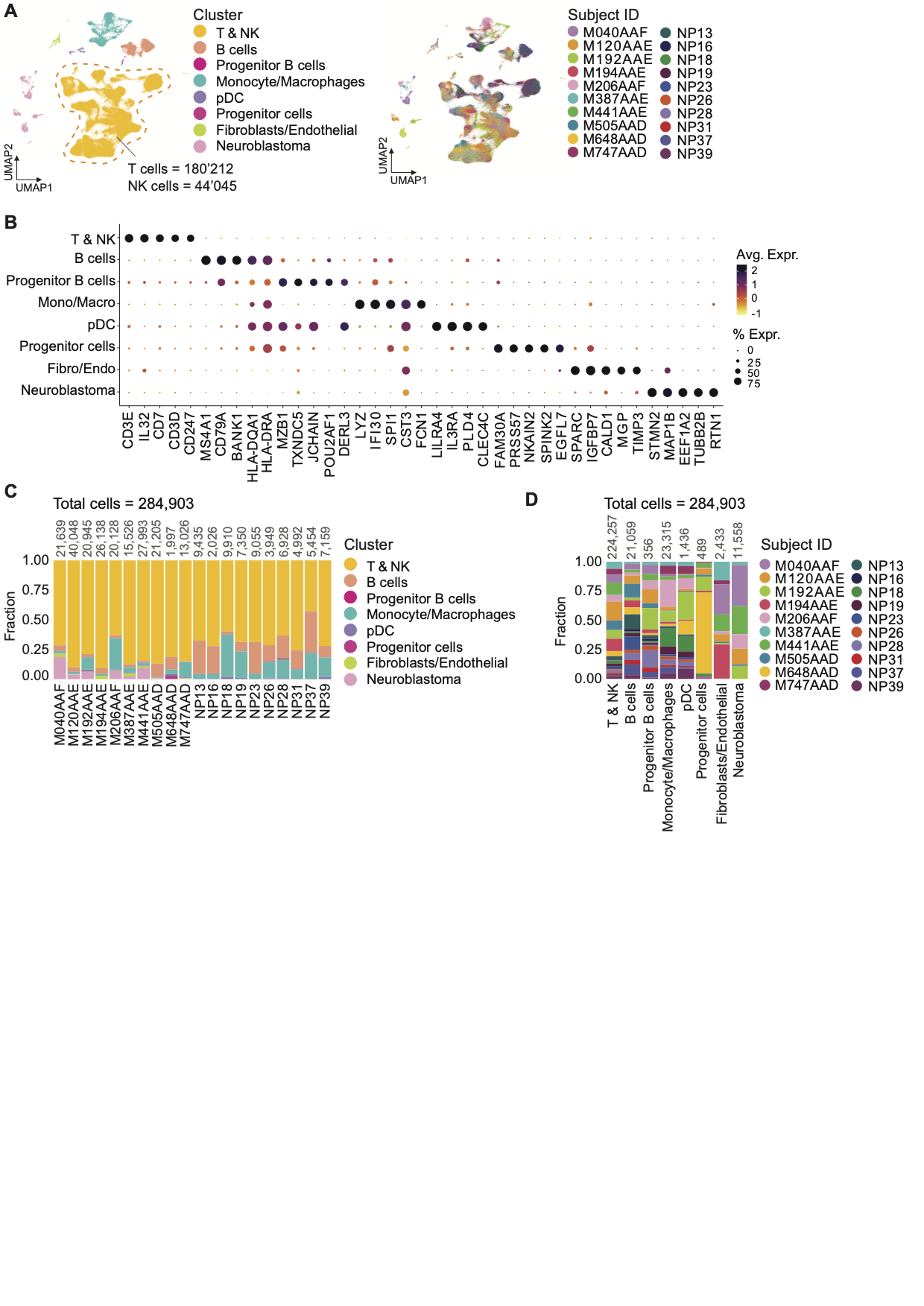

### Figure S2

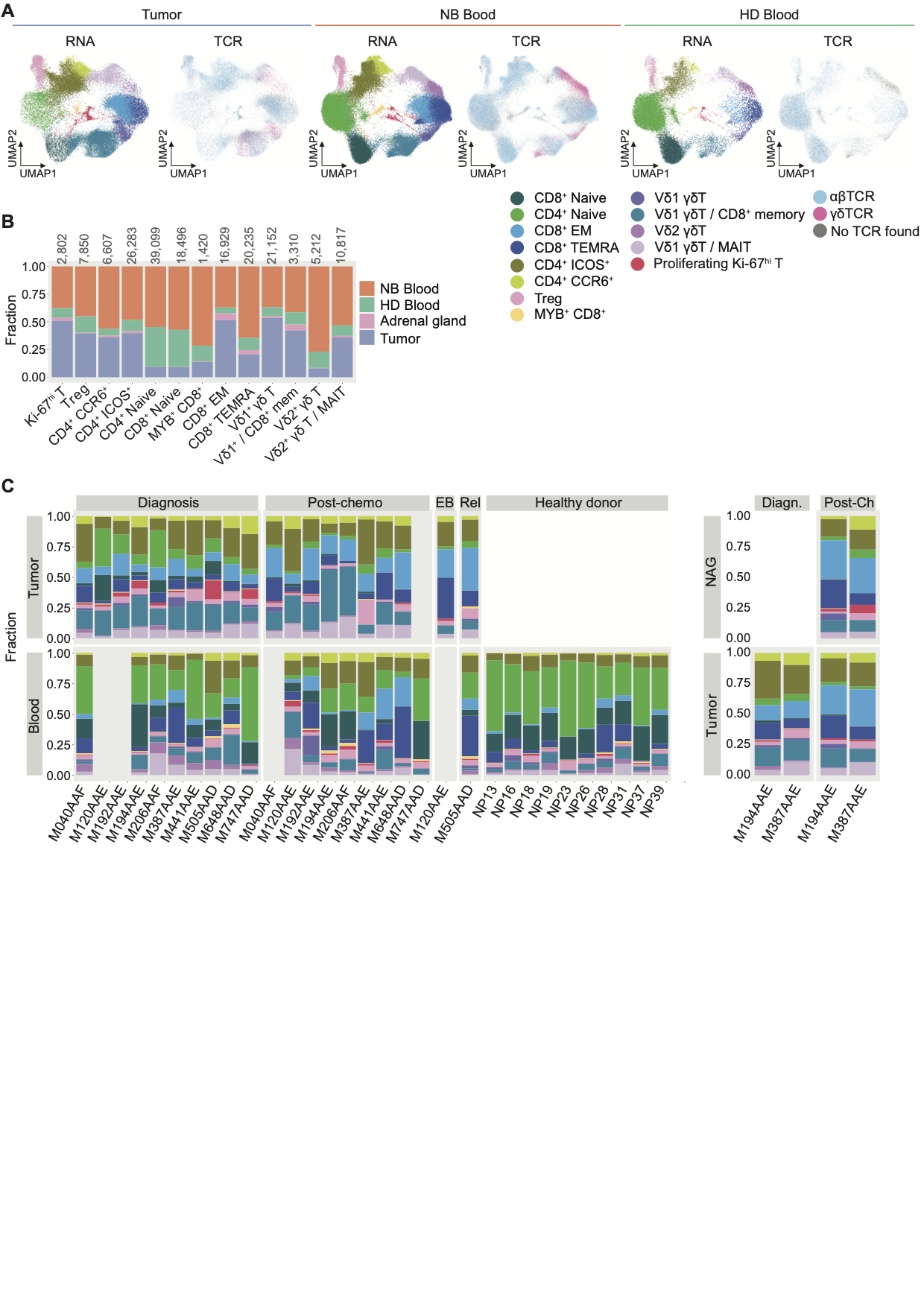

### Figure S3

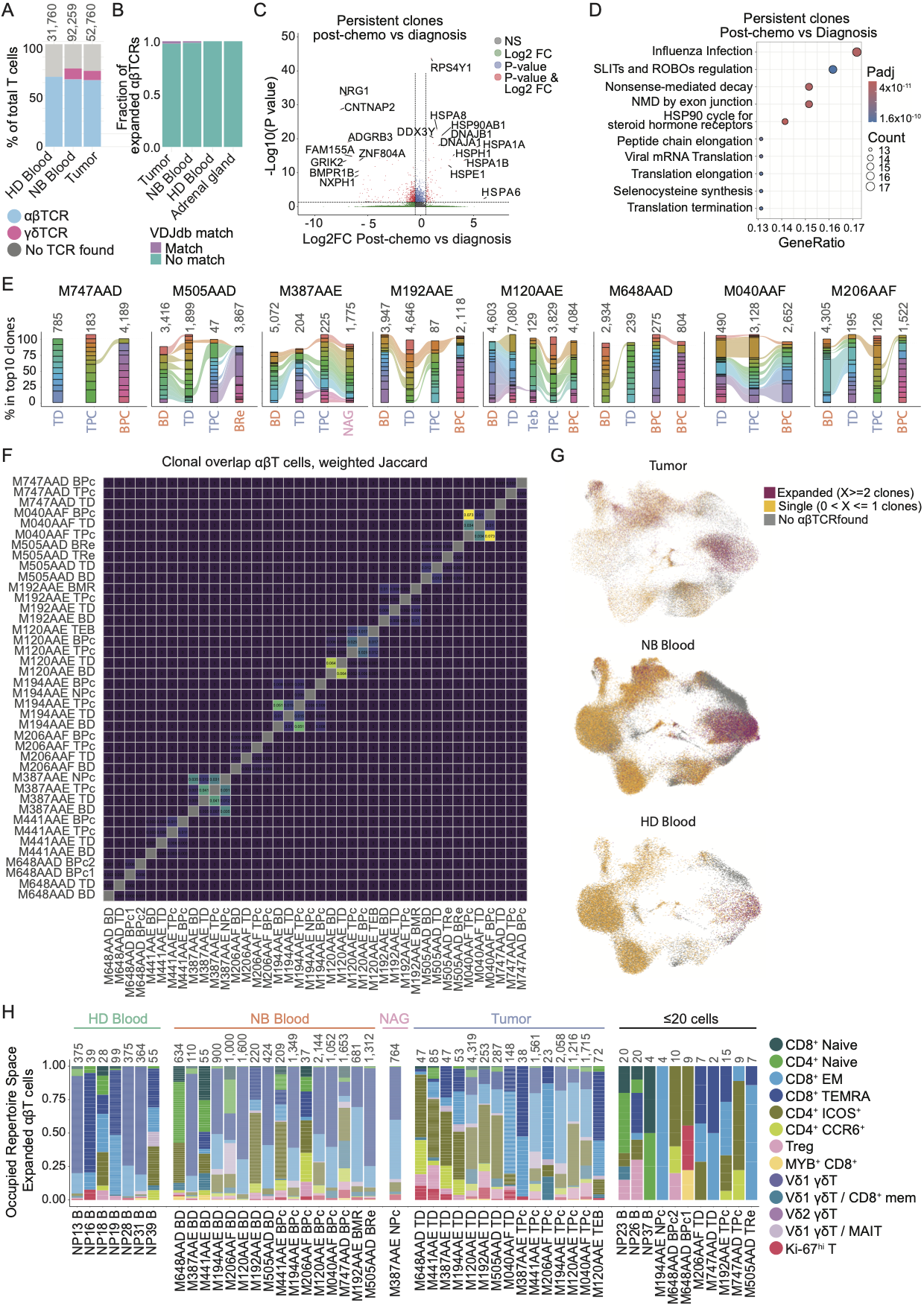

### Figure S4

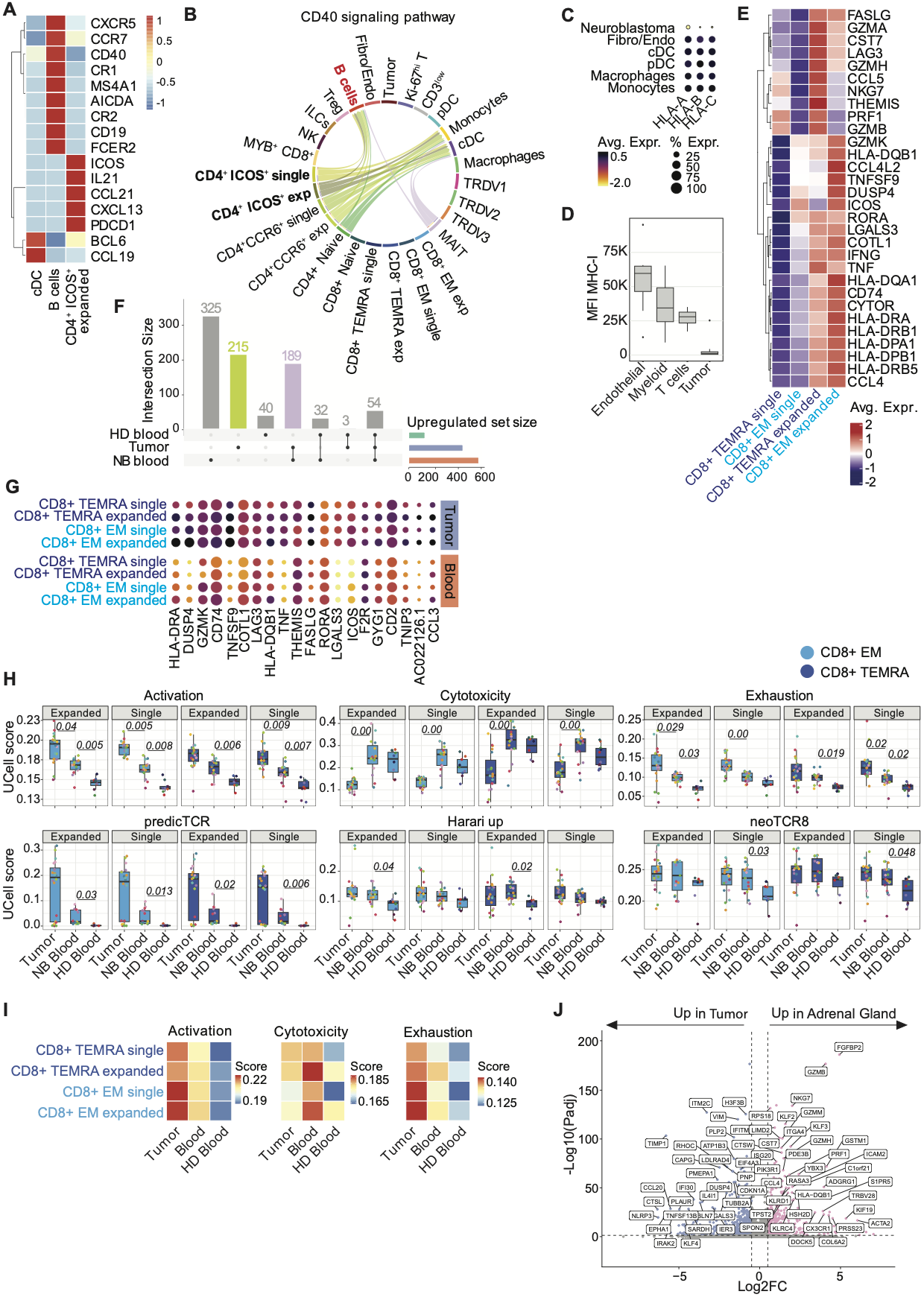

### Figure S5

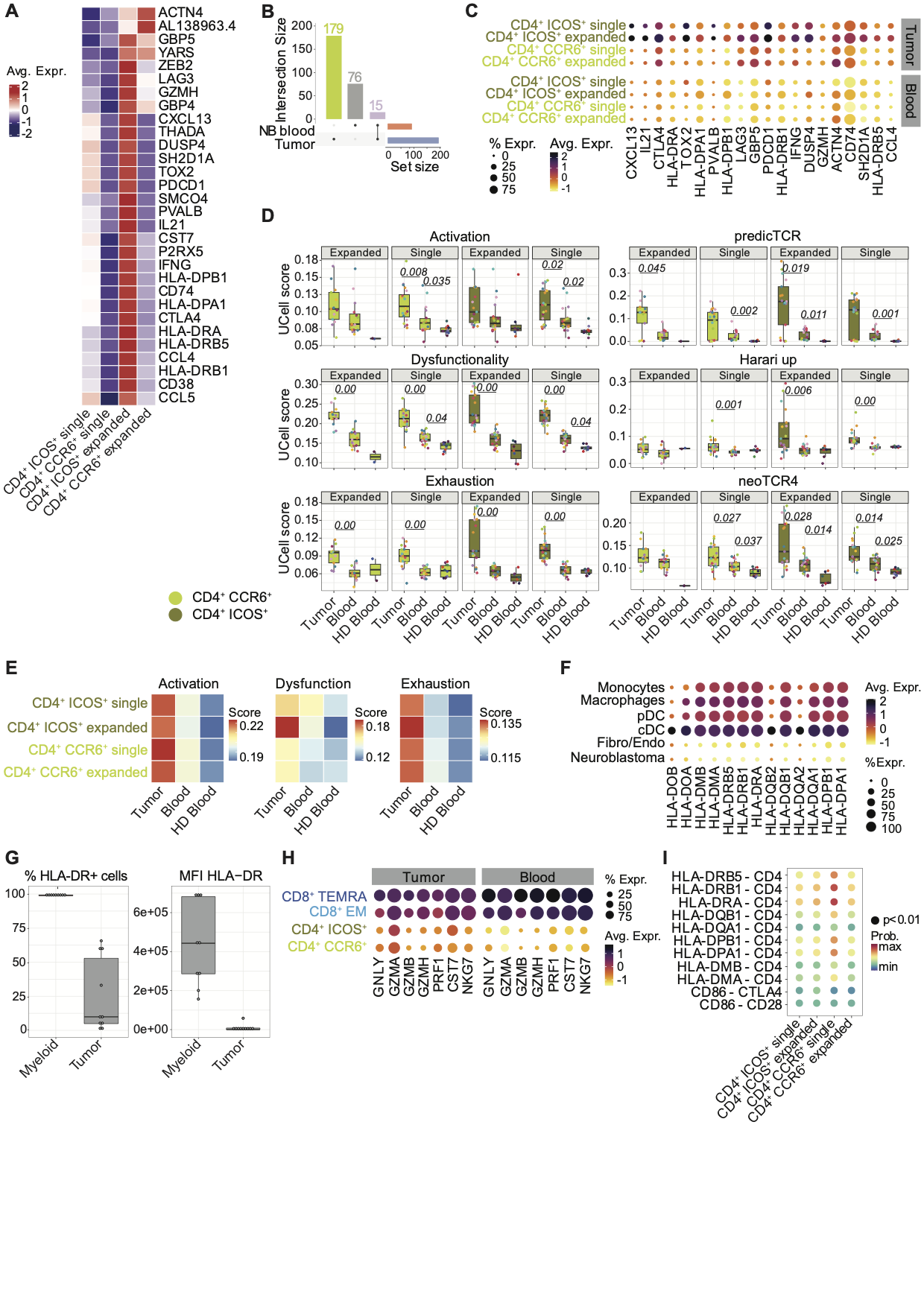

### Figure S6

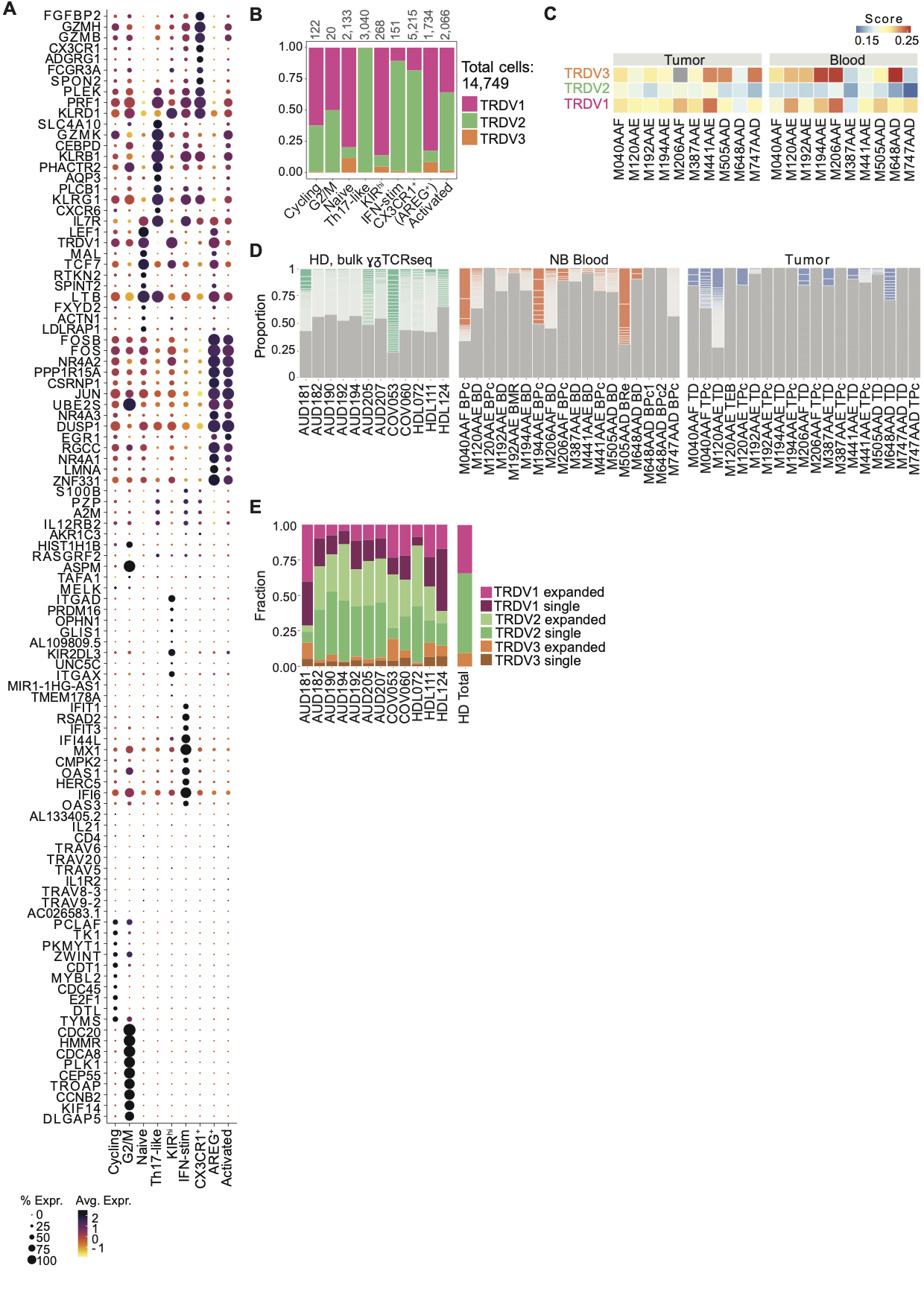

### Figure S7

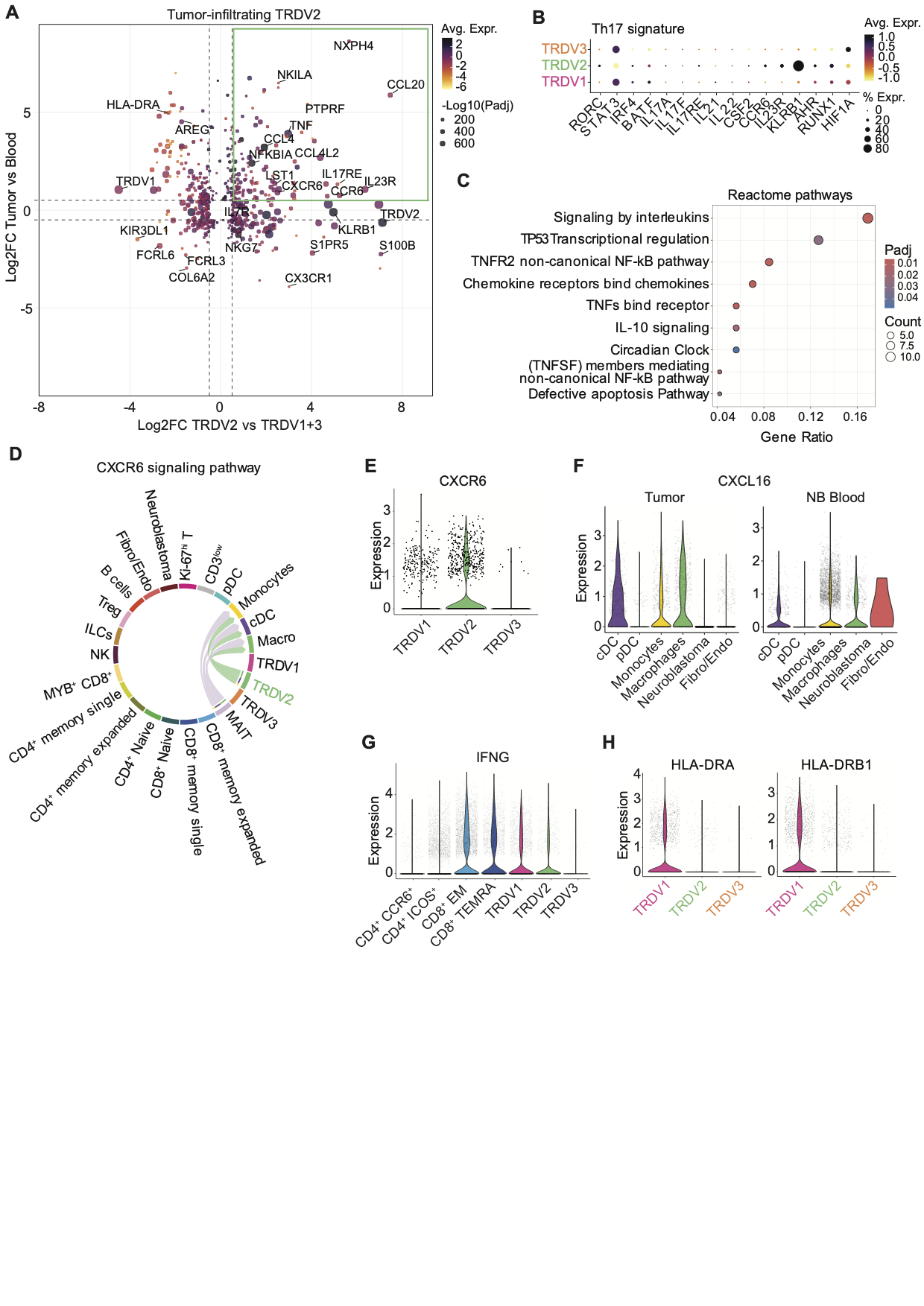

### Figure S8

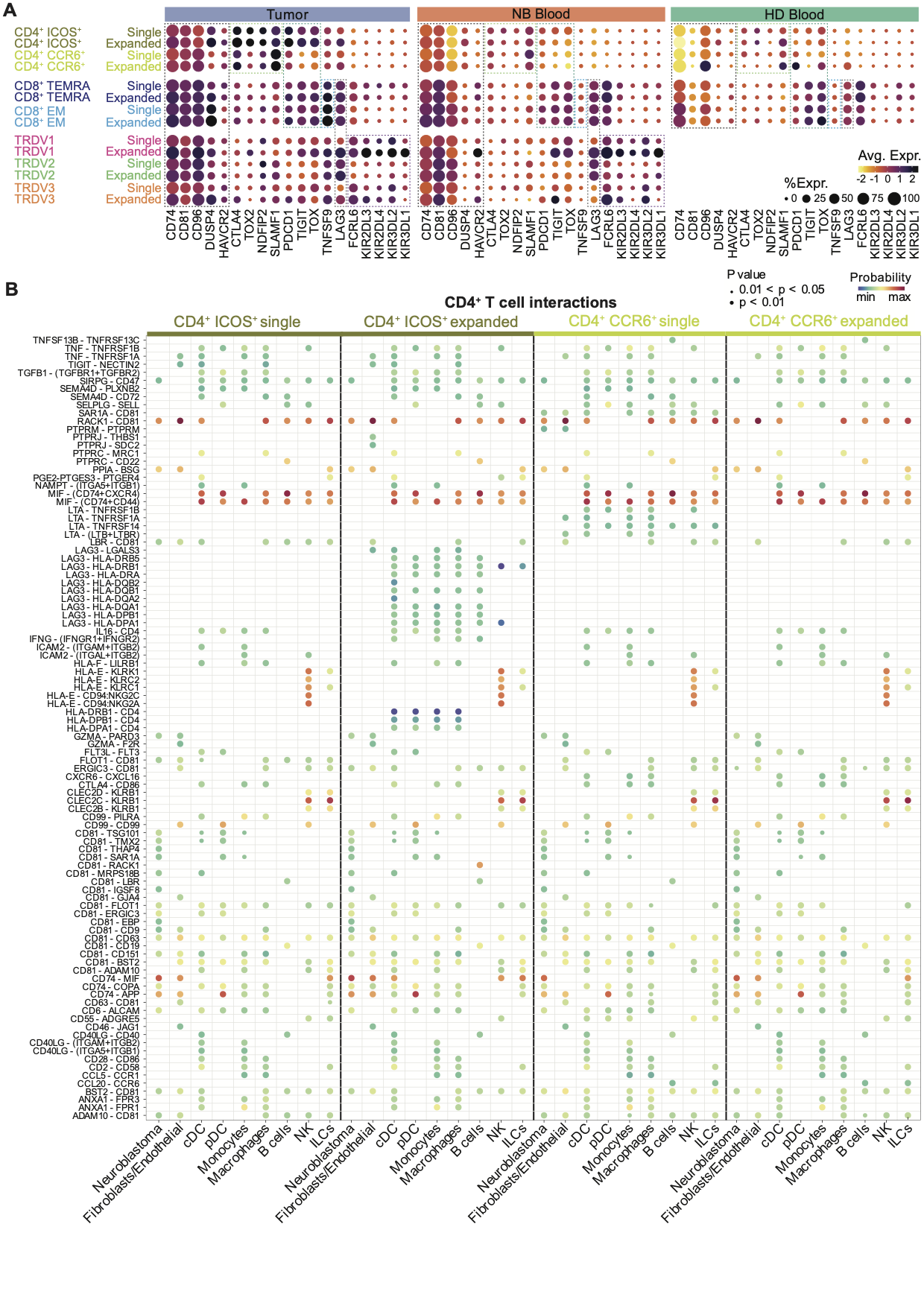

### Figure S9

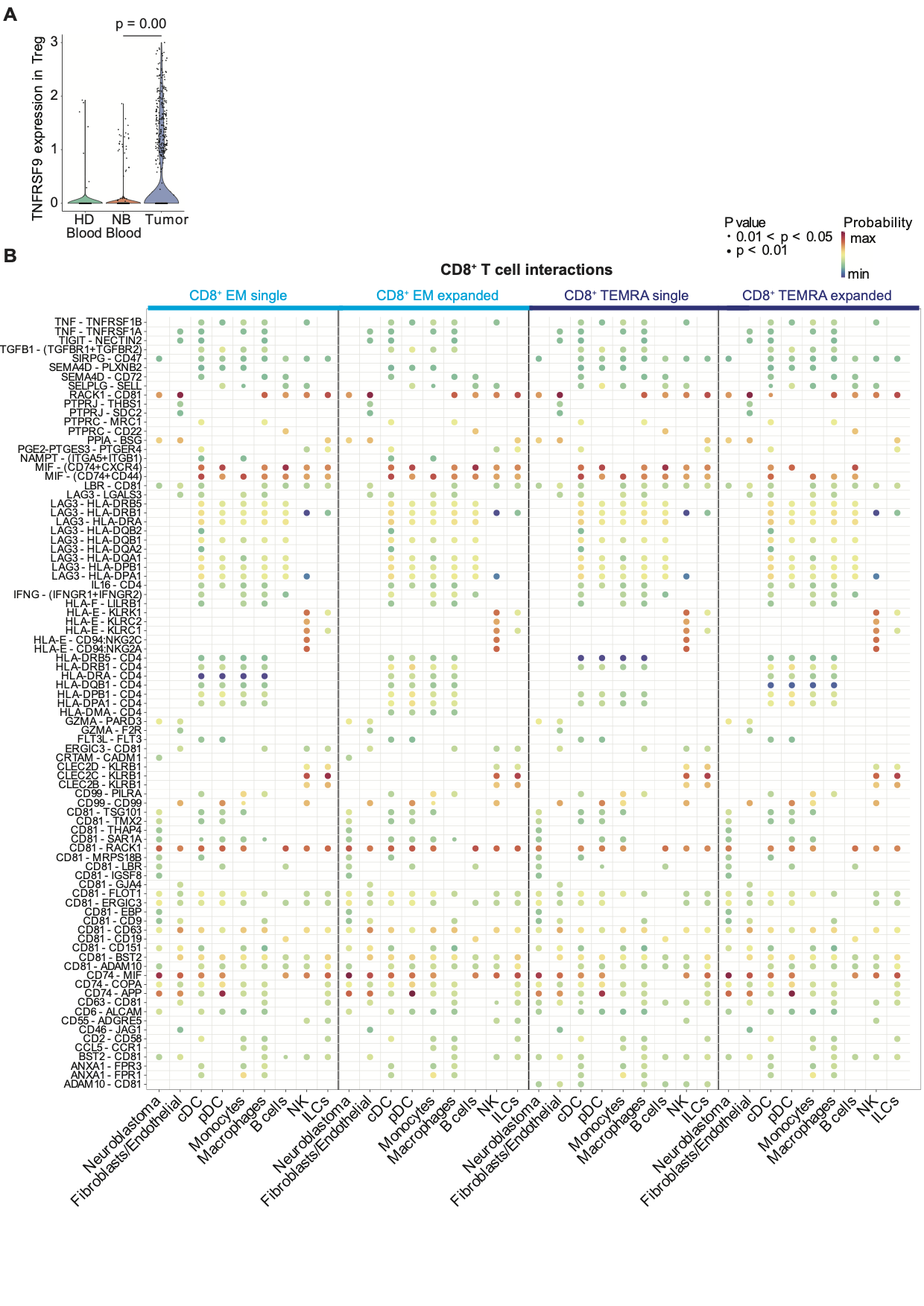

### Figure S10

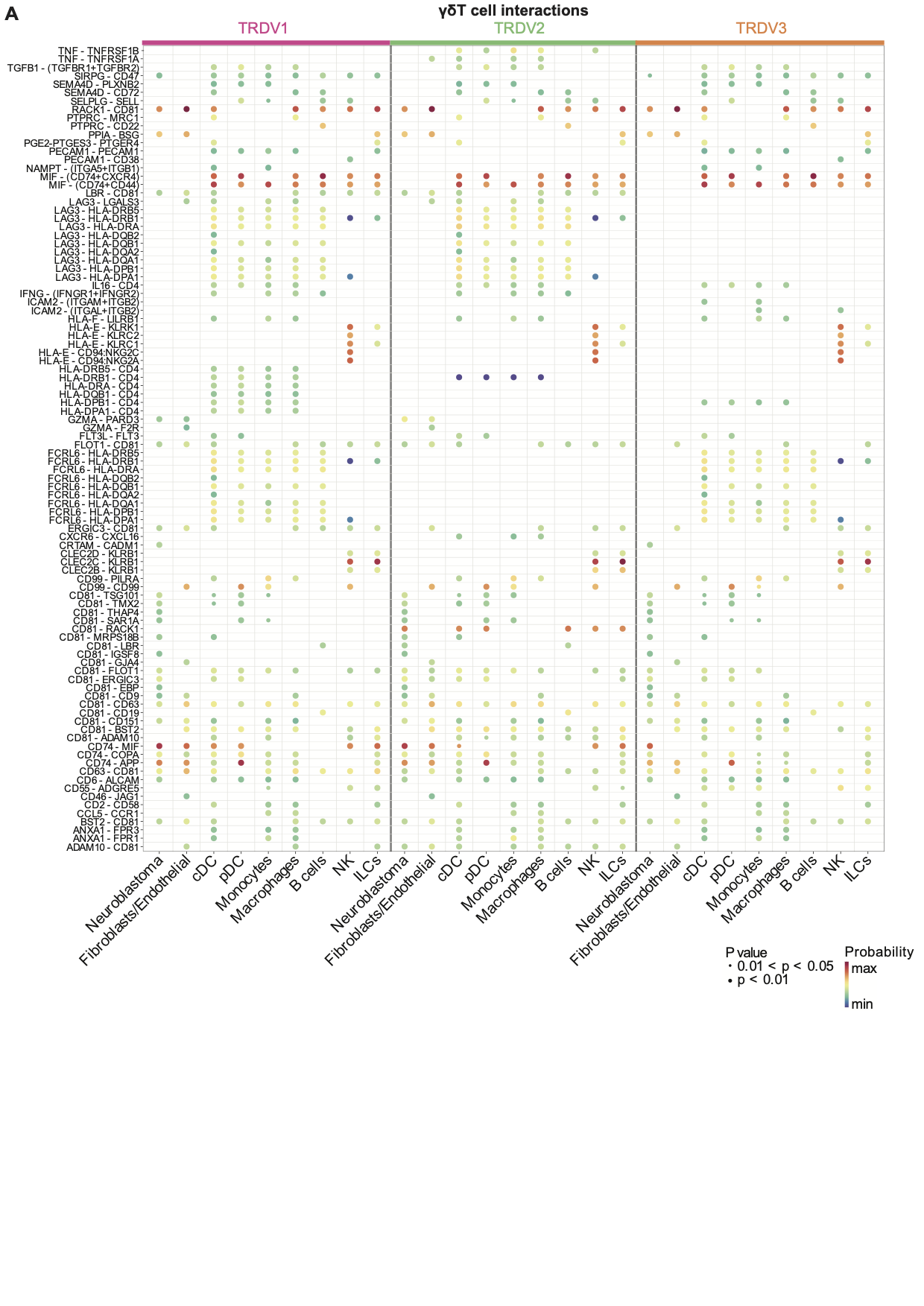

### Supplementary methods

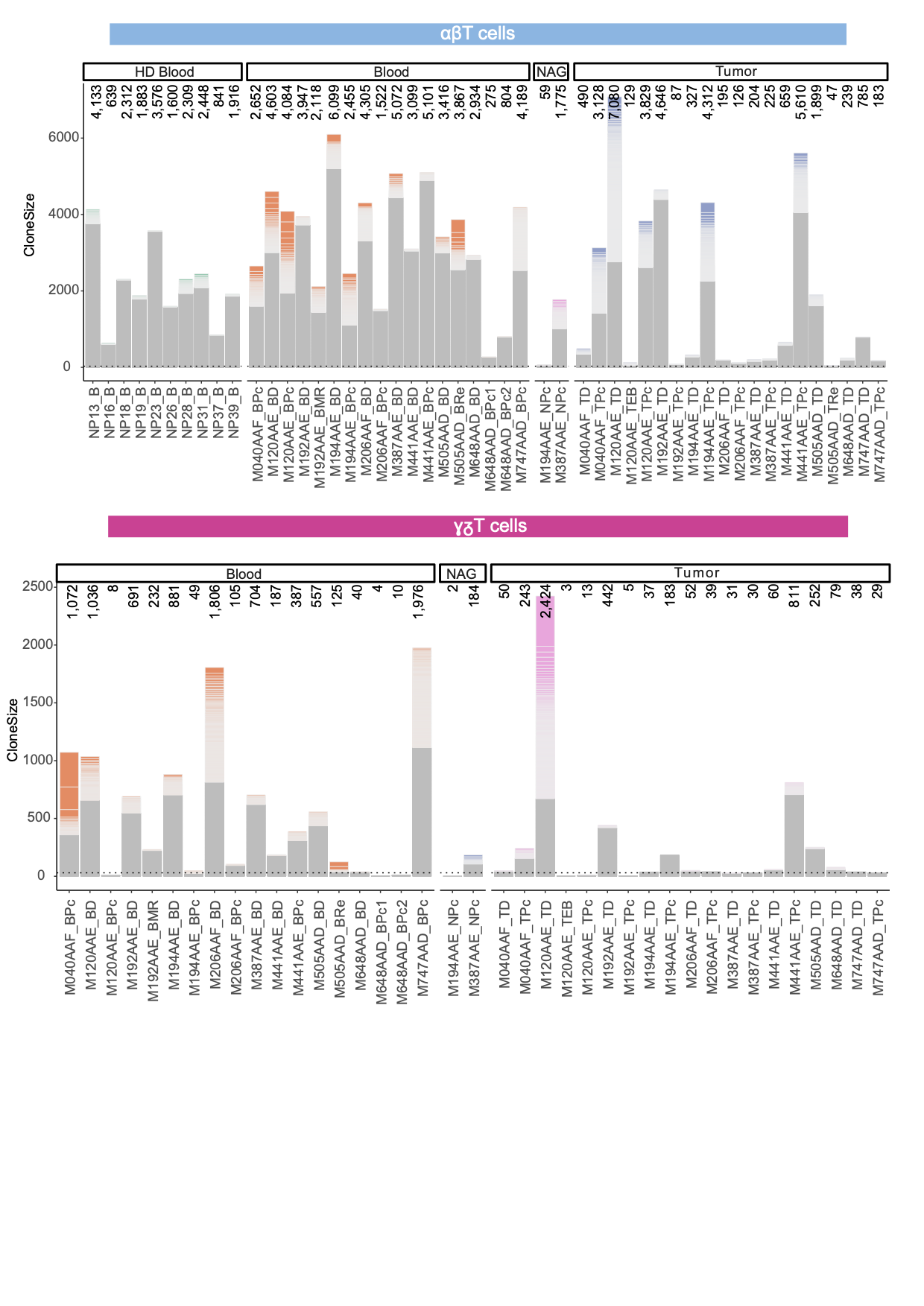
